# Parallel and lineage-dependent mitonuclear evolution in threespine stickleback

**DOI:** 10.64898/2026.11.02.763752

**Authors:** Changyue Yu, Daniel I. Bolnick, Jesse N. Weber

## Abstract

Mitochondrial function depends on interactions between proteins and RNAs encoded by the mitochondrial and nuclear genomes. Repeated adaptation to similar environments may favor parallel changes in nuclear genes supporting mitochondrial function, while maintaining compatibility with different mitochondrial backgrounds may favor population-specific responses. Threespine stickleback (*Gasterosteus aculeatus*) repeatedly colonized freshwater habitats from ancestral marine populations carrying standing genetic variation. We tested whether these colonizations resulted in parallel evolution of nuclear oxidative phosphorylation (OXPHOS) genes and whether these responses coincided with evidence of mitonuclear evolutionary coupling. Several divergent mitochondrial lineages were distributed across populations with limited geographic correspondence. Mitochondrial ribosomal proteins and aminoacyl-tRNA synthetases exhibited elevated nonsynonymous divergence and polymorphism relative to their cytosolic counterparts. We also identified positive evolutionary rate covariation between mitochondrial and nuclear genes that comprise OXPHOS Complex IV, suggesting that evolutionary coupling is concentrated in particular functional modules. Many nuclear OXPHOS genes displayed large allele-frequency changes that were parallel across freshwater populations in Alaska and British Columbia, while the magnitude of differentiation differed between regions. Mitochondrial lineage was associated with a smaller, locus-specific component of nuclear OXPHOS variation, concentrated in Complex I genes. Notably, the genes with large freshwater-associated evolutionary shifts were largely distinct from those displaying mitonuclear rate covariation. In this context, the gene *cox4i1* emerges as a candidate connecting repeated freshwater adaptation with molecular variation at the mitonuclear interface. Together, these findings suggest that freshwater adaptation drives broadly parallel nuclear OXPHOS evolution across heterogeneous mitochondrial backgrounds, while evidence of mitonuclear evolutionary coupling is more localized.

## Introduction

Mitochondrial and nuclear genomes are deeply integrated components of the eukaryotic cell (Poyton and McEwen 1996). Although mitochondria retain a small, independently inherited genome, most proteins required for mitochondrial function are encoded in the nucleus and imported post-translationally. Core processes such as oxidative phosphorylation (OXPHOS), mitochondrial translation, and mitochondrial genome maintenance therefore depend on coordinated interactions between products of the two genomes (Gray et al. 1999; Wallace 2007; Calvo and Mootha 2010). OXPHOS couples electron transport to ATP synthesis through five protein complexes, four of which contain subunits encoded by both the mitochondrial and nuclear genomes. This functional interdependence creates an evolutionary puzzle. In many animals, mitochondrial DNA (mtDNA) combines elevated mutation rates with reduced effective population size and limited recombination, potentially increasing the accumulation of mildly deleterious mutations; yet mitochondrial function is generally maintained across populations and species (Lynch and Blanchard 1998). Moreover, because mitochondrial function underlies energy production and metabolic performance, divergence in mtDNA may do more than require the conservation of existing function. It may alter the fitness effects of nuclear alleles and thereby influence the adaptive trajectories available to populations.

A widely invoked explanation for sustained mitonuclear compatibility is compensatory coevolution. Mutations in one genome, often the rapidly evolving mitochondrial genome, can impose selection for compatible changes in the other (Rand et al. 2004). Under the nuclear compensation hypothesis, destabilizing or mildly deleterious mitochondrial mutations favor compensatory evolution in nuclear genes whose products interact with mitochondrial-encoded proteins or RNAs (Dowling et al. 2008; Osada and Akashi 2012). The fitness effects of nuclear alleles can depend on their mitochondrial background, with incompatible combinations impairing respiratory function and survival (Moran et al. 2024). Beyond their implications for hybrid incompatibility and reproductive isolation, mitonuclear interactions may also shape metabolic performance and influence the genetic responses available during adaptation to different environmental conditions (Bettinazzi et al. 2024; Kayhani et al. 2026). Such background dependence raises the possibility that populations carrying divergent mitochondrial backgrounds may favor different nuclear variants during adaptation to similar environments (Hill 2015; Hill et al. 2019).

Mitonuclear coevolution can involve gene products contributing to several mitochondrial functions. These include nuclear-encoded OXPHOS subunits that assemble with mitochondrially encoded respiratory-chain components, mitochondrial aminoacyl-tRNA synthetases that charge mitochondrial tRNAs, and mitoribosomal proteins that interact with mitochondrially encoded rRNAs. Physical contact may strengthen evolutionary coupling, although its contribution varies among taxa and is not consistently significant (Yan et al. 2019; Piccinini et al. 2021; Weaver et al. 2022). Comparative studies have identified elevated evolutionary rates in mitochondrial-associated nuclear genes and coordinated evolutionary rates between mitochondrial and nuclear components, consistent with mitonuclear coevolution (Barreto et al. 2018; Yan et al. 2019; Weaver et al. 2022). Comparisons of mitochondrial translation proteins with their functionally analogous cytosolic counterparts provide an additional means of evaluating whether evolutionary differences are associated with mitochondrial function. However, accelerated evolution alone cannot distinguish compensatory coevolution from relaxed selective constraint, population history, or other selective forces. Much of the comparative evidence also comes from deeply diverged lineages, leaving the relationship between mitonuclear interactions and recent, within-species adaptation less well understood.

Ecological selection provides another potential driver of mitochondrial and nuclear evolution. Differences in temperature and oxygen availability can alter energetic demands and impose selection on mitochondrial pathways (Ballard and Melvin 2010; Wilson et al. 2012). Biotic conditions may also contribute: mitochondrial reactive oxygen species participate in innate immune signaling and antimicrobial responses (West et al. 2011). Because parasite-mediated selection can favor divergent immune responses among stickleback populations (Weber et al. 2022), variation in parasitism could also indirectly impose selection on mitochondrial function. Repeated exposure to similar environments may favor parallel evolutionary responses in nuclear genes that support mitochondrial function. However, whether these responses consistently involve the same genes, and whether they coincide with signatures of mitonuclear coevolution, remain unresolved. Similar ecological transitions need not produce identical genetic outcomes. If the fitness effects of nuclear alleles depend on mitochondrial background, shared ecological pressures may favor broadly parallel responses, while mitochondrial background may influence which nuclear variants are favored in individual populations. Examining whether environmentally associated parallelism and mitochondrial-lineage-associated nuclear variation involve the same genes is therefore important for understanding how ecological adaptation and mitonuclear interactions jointly contribute to adaptive diversification.

Threespine stickleback (*Gasterosteus aculeatus*) provides a powerful system in which to test this framework. Following Pleistocene deglaciation, marine stickleback repeatedly colonized a large number of freshwater habitats across the Northern Hemisphere, generating independent episodes of ecological and genomic divergence (Bell and Foster 1994; Jones et al. 2012). These colonizations exposed populations to substantial changes in salinity, temperature, oxygen availability, and resource conditions, all of which may reshape selection on energy metabolism (Pörtner and Farrell 2008; Aguirre et al. 2022). At the same time, stickleback populations harbor deeply divergent mitochondrial haplotypes, including variation in protein-coding genes that encode core OXPHOS subunits (Beck et al. 2022). This variation likely reflects the presence of multiple ancestral mitochondrial lineages in marine populations and their differential sampling during the colonization of freshwater drainages (Thompson et al. 1997). Independent freshwater populations can therefore experience broadly similar ecological transitions while carrying distinct mitochondrial backgrounds, providing an opportunity to examine shared ecological responses alongside mitochondrial-lineage-associated nuclear variation.

Here, we investigate how environmental change and mitochondrial genetic background jointly shape mitonuclear evolution during the adaptive radiation of threespine stickleback. We first characterize mitochondrial variation among populations and compare molecular divergence and within-population diversity across mitochondrial-associated and cytosolic gene classes. We then test whether mitochondrial- and nuclear-encoded genes show coordinated evolutionary rates and whether repeated freshwater colonization is associated with parallel allele-frequency shifts in nuclear OXPHOS genes. To distinguish broad spatial patterns from mitochondrial effects, we separately test associations of nuclear OXPHOS variants with geographic coordinates and mitochondrial lineage and compare nuclear evolutionary trajectories among populations sharing or differing in mitochondrial background. Finally, we identify recurrent freshwater-associated amino-acid substitutions and evaluate their predicted functional and structural consequences. Together, these analyses assess broad ecological parallelism alongside more localized signatures of mitonuclear evolutionary coupling.

## Results

### Mitochondrial genetic variation across and within populations

To characterize mitochondrial genetic variation among stickleback populations, we analyzed previously published whole-genome Pool-seq data from 27 stickleback populations spread across Vancouver Island, BC, Canada and Alaska, USA, including two ancestral marine populations and 25 freshwater populations, five of which were recently colonized (Fig. S1; Table S1; 46–200 individuals per population; Weber et al. 2022). A median-joining haplotype network based on population-consensus mitogenomes revealed several differentiated mitochondrial lineages connected through a central group containing the two anadromous marine populations (Fig. 1A). We also reconstructed a maximum-likelihood phylogeny from the population-consensus mitochondrial sequences, using Japan Sea stickleback (*G. nipponicus*) as the outgroup (Fig. 1B). Branch support varied across the mitochondrial phylogeny, with lower support for many shallow relationships (seven of the 25 internal branches met both support thresholds: SH-aLRT ≥ 80% and ultrafast bootstrap ≥ 95%). The network and phylogeny recovered several similar population groupings but differed in some weakly supported fine-scale relationships, which we did not interpret further.

**Figure 1.**
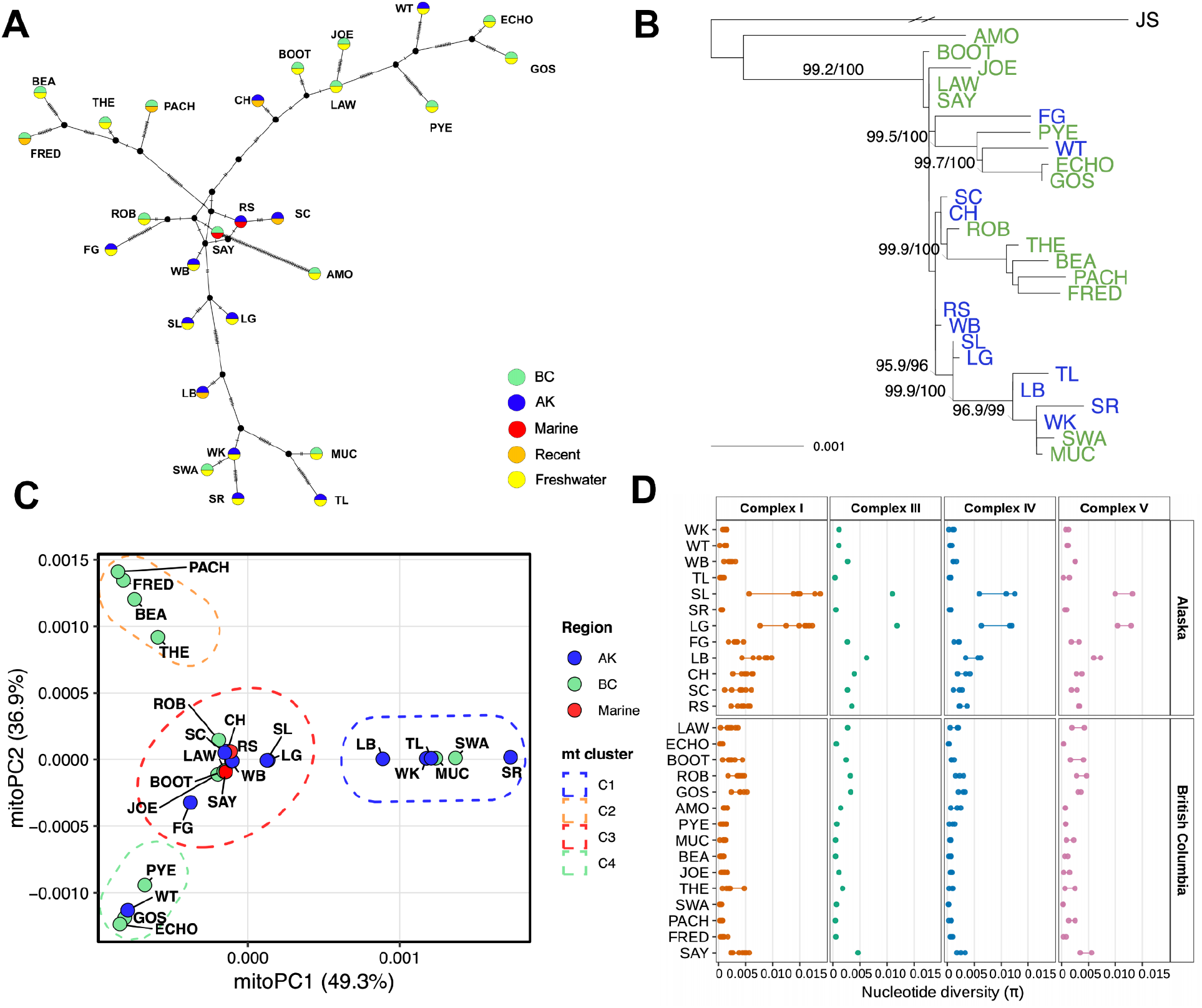
Mitochondrial genetic structure of 27 stickleback populations. (A) Haplotype network based on population level consensus mtDNA sequences using Median Joining methods. Each node represents a unique haplotype. Tick marks represent mutational steps. (B) Maximum-likelihood phylogeny constructed from population-consensus mitochondrial sequences after excluding the control region (D-loop), with Japan Sea stickleback (*Gasterosteus nipponicus*) as the outgroup. Branch lengths represent substitutions per site, except that the long branch leading to the outgroup was truncated for visualization and is marked by a double slash. Node labels show SH-aLRT/ultrafast-bootstrap support and are displayed when SH-aLRT ≥ 80% and ultrafast bootstrap ≥ 95%. Tip-label colors indicate geographic region. (C) Mitochondrial PCoA analysis based on mitochondrial phylogenetic tree (excluding the AMO population, which displayed extreme divergence from other populations and thus skewed PCoA patterns). (D) Gene-level mitochondrial nucleotide diversity (π) across populations, partitioned by geographic region and OXPHOS complex. Points represent individual mitochondrial genes.

AMO was recovered as a sister to the remaining 26 northeastern Pacific populations with strong support (SH-aLRT = 99.2%; ultrafast bootstrap = 100%). It also had the longest terminal branch among the focal populations (0.00240 substitutions per site), approximately 2.3 times the length of the next-longest terminal branch. Because this divergent lineage was represented by a single population and strongly influenced distance-based ordination, AMO was retained in the phylogenetic and nucleotide-diversity analyses but excluded from the PCoA and subsequent analyses requiring categorical mitochondrial-lineage assignments. After excluding AMO, PCoA and *k*-means clustering summarized the remaining mitochondrial variation into four clusters that were broadly consistent with the major groupings in the haplotype network (Fig. 1C). Mitochondrial cluster membership was not significantly associated with watershed grouping after controlling for geographic region (within-region permutation test, *P* = 0.263).

Watershed-based prediction also did not significantly improve upon region-only prediction in leave-one-population-out cross-validation (35.1% vs. 26.3% accuracy; permutation *P* = 0.274). We next evaluated within-population mtDNA diversity (π) for each population. Among OXPHOS complexes, Complex I genes (ND1, ND2, ND4, ND5) consistently exhibited the highest nucleotide diversity across populations, followed by Complex V (ATP6, ATP8) and Complex IV (COX1–3), whereas Complex III (CYTB) showed comparatively lower diversity (Fig. 1D). Complex I had the highest diversity in 12 out of 27 populations (44.4%), significantly exceeding the expectation under random assignment (binomial test, *P* = 0.022). Nucleotide diversity did not vary significantly among populations (Kruskal–Wallis test, *P* = 0.463), but the highest values were observed in Alaskan freshwater populations. In particular, Long Lake (LG) and Spirit Lake (SL) exhibited approximately three-fold higher π than most other populations. Although we could not reconstruct individual-level haplotypes from Pool-seq data, the high π in LG and SL suggests that divergent haplotypes may be segregating in these populations.

### Ortholog identification and gene-set curation

Although the threespine stickleback genome is relatively well annotated, orthology has not been clearly assigned to all nuclear genes with mitochondrial function. We retained the genome-wide classification of nuclear genes as direct n-mt (encode products with documented physical or molecular interactions with mt proteins or RNAs), indirect n-mt (encode mt localized products without known direct interactions), or non-n-mt (no mitochondrial association). We also curated gene sets representing OXPHOS structural subunits and assembly factors, mitochondrial aminoacyl-tRNA synthetases, and mitochondrial ribosomal proteins, together with functionally analogous cytosolic comparison groups. OXPHOS structural subunits were further classified as core (bacterial homologs; direct catalytic, electron-transfer, or proton-translocation roles) or noncore (later-derived accessory subunits). Detailed definitions, orthology criteria, and gene membership are provided in the Materials and Methods and Table S2.

### Molecular evolution varies among mitochondrial functional modules

To characterize molecular evolution across mitochondrial-associated gene classes, we compared divergence estimated from quality-filtered population-consensus coding sequences (d_N_, d_S_, and d_N_/d_S_) and within-population nucleotide diversity (π_N_, π_S_, and π_N_/π_S_) (Tables S3 and S4; Fig. 2 and Figs. S2–S5). Across OXPHOS-related comparisons, differences in d_N_/d_S_ were limited to specific gene classes. Mitochondrial OXPHOS genes (mt OXPHOS), nuclear-encoded OXPHOS structural subunits (nu OXPHOS), and nuclear OXPHOS assembly factors (nu assembly factors) did not differ significantly in d_N_ (Fig. 2A). Mt OXPHOS genes had significantly higher d_S_ than both nuclear classes and higher d_N_/d_S_ than nu OXPHOS genes, which is consistent with higher mutation rates in mitochondrial genomes and stronger purifying selection operating on nu OXPHOS genes. However, several nu OXPHOS genes (*cox4i1* and *ndufs7*) did show evidence of positive selection (i.e., d_N_/d_S_ > 1; Fig. 2A; Table S3). Across respiratory-chain complexes, significant gene-class differences in d_S_ were observed in Complex I, whereas no complex showed significant differences in d_N_ or d_N_/d_S_ (Fig. S3A). Within populations, π_N_ and π_S_ differed among the broad OXPHOS gene classes, but π_N_/π_S_ did not (Fig. S2A). Across respiratory-chain complexes, differences in π_N_ and π_S_ were concentrated primarily in Complex I, with an additional π_N_ difference in Complex II; however, no complex showed significant gene-class differences in π_N_/π_S_ (Fig. S3B). Together, these comparisons suggest that selective constraints may differ among specific OXPHOS gene classes.

**Figure 2.**
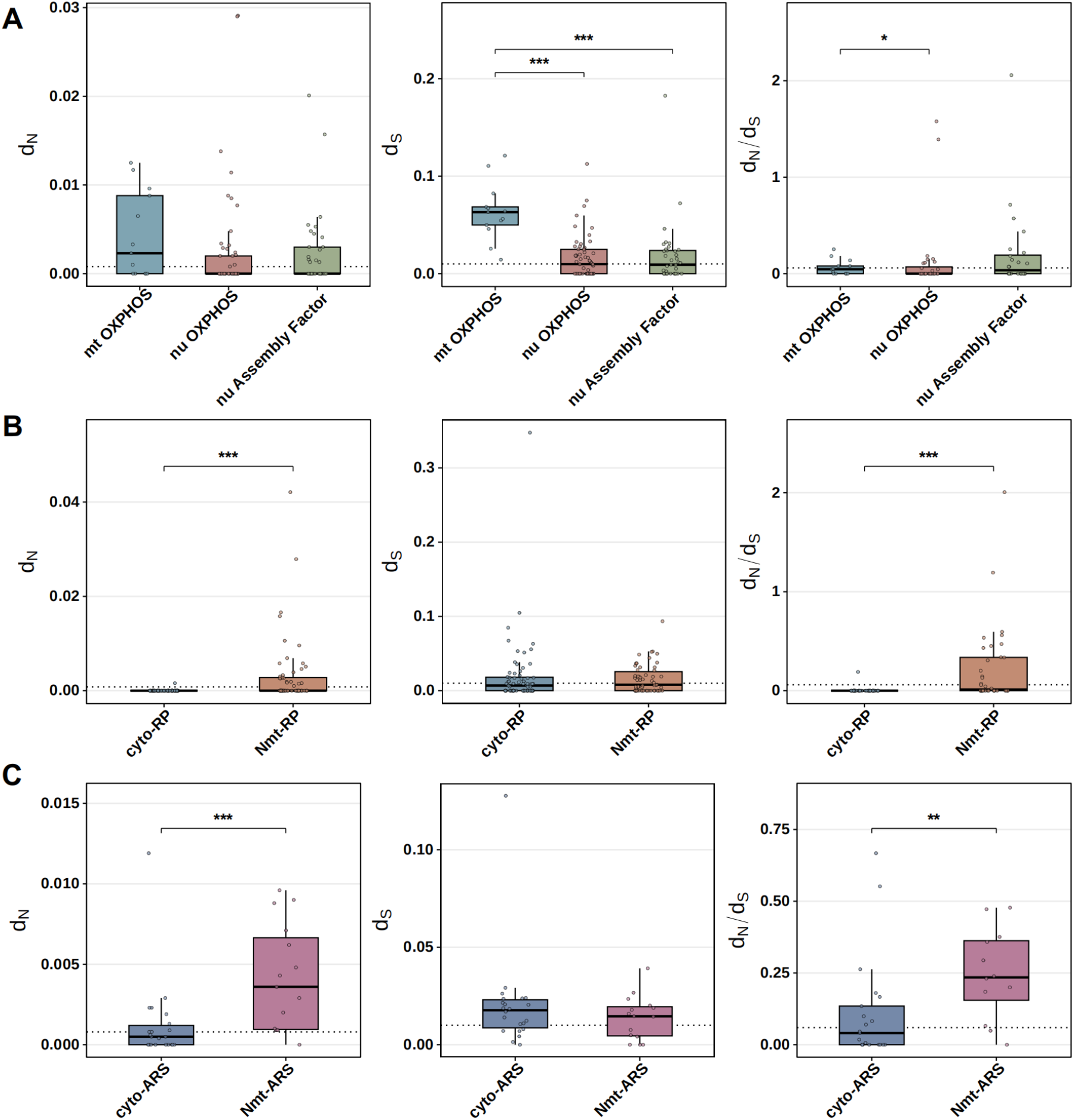
Divergence patterns across OXPHOS, ribosomal protein, and aminoacyl-tRNA synthetase genes reveal differences in evolutionary constraint. Distributions of d_N_, d_S_, and d_N_/d_S_ are shown for (A) mitochondrial OXPHOS genes (mt OXPHOS), nuclear OXPHOS subunit genes (nu OXPHOS), and nuclear OXPHOS assembly-factor genes; (B) cytosolic and nuclear-encoded mitochondrial ribosomal proteins (RPs); and (C) cytosolic and nuclear-encoded mitochondrial aminoacyl-tRNA synthetases (ARSs). Boxes indicate medians and interquartile ranges; whiskers extend to the most extreme values within 1.5 times the interquartile range. Dotted horizontal lines indicate genome-wide nuclear medians. Pairwise differences were evaluated using two-sided Wilcoxon rank-sum tests with Benjamini–Hochberg correction. Asterisks denote adjusted significance levels *P* < 0.05 (*), *P* < 0.01 (**), *P* < 0.001 (***).

Mitochondrial ribosomal proteins (Nmt-RPs) and mitochondrial aminoacyl-tRNA synthetases (Nmt-ARSs) showed clear differences in molecular evolution compared to their cytosolic counterparts. Nmt-RPs had significantly higher d_N_ and d_N_/d_S_ than cytosolic ribosomal proteins (Cyto-RPs), with no corresponding difference in d_S_ (Fig. 2B). Within populations, Nmt-RPs also had higher π_N_ and π_N_/π_S_ than Cyto-RPs, whereas π_S_ did not differ between the two classes (Fig. S2B). Cyto-RPs generally fell below the corresponding genome-wide medians for d_N_ and π_N_, whereas Nmt-RPs were closer to or above the genome-wide background. An even stronger pattern was observed for mitochondrial aminoacyl-tRNA synthetases (Nmt-ARSs; Fig. 2C). Nmt-ARSs had significantly higher d_N_ and d_N_/d_S_ than cytosolic aminoacyl-tRNA synthetases (Cyto-ARSs), and their median values for both measures exceeded the corresponding genome-wide nuclear medians. Within populations, Nmt-ARSs also had higher π_N_ and π_N_/π_S_ but lower π_S_ than Cyto-ARSs (Fig. S2C). Thus, the elevated d_N_/d_S_ and π_N_/π_S_ of Nmt-RPs and Nmt-ARSs were not accompanied by increased synonymous divergence or diversity, consistent with differences in selective constraint.

Nuclear core subunits had significantly higher d_N_ and d_N_/d_S_ than nuclear noncore subunits, whereas π_N_/π_S_ did not differ significantly (Fig. S4). In contrast, direct n-mt, indirect n-mt, and non-n-mt genes did not differ significantly in d_N_, d_S_, d_N_/d_S_, or π_N_; however, indirect n-mt genes had significantly higher π_S_ and lower π_N_/π_S_ than non-n-mt genes (Fig. S5). These results indicate that patterns of divergence and nucleotide diversity varied among mitochondrial-associated functional modules, rather than showing a uniform shift across all OXPHOS or nuclear-mitochondrial genes.

### Evolutionary rate covariation reveals complex-specific mitonuclear coupling

To test whether mitochondrial and nuclear genes showed evolutionary-rate covariation across lineages, we quantified the Spearman correlation between branch-specific relative evolutionary rates of the composite set of 13 mitochondrial protein-coding genes and individual nuclear genes across different functional sets (Fig. 3; Table S5). Evolutionary-rate covariation was not broadly elevated among nuclear genes associated with mitochondrial function. The bootstrap confidence intervals for direct and indirect n-mt genes, nu OXPHOS subunits, OXPHOS assembly factors, Nmt-RPs and Nmt-ARSs all overlapped zero (Fig. 3A).

**Figure 3.**
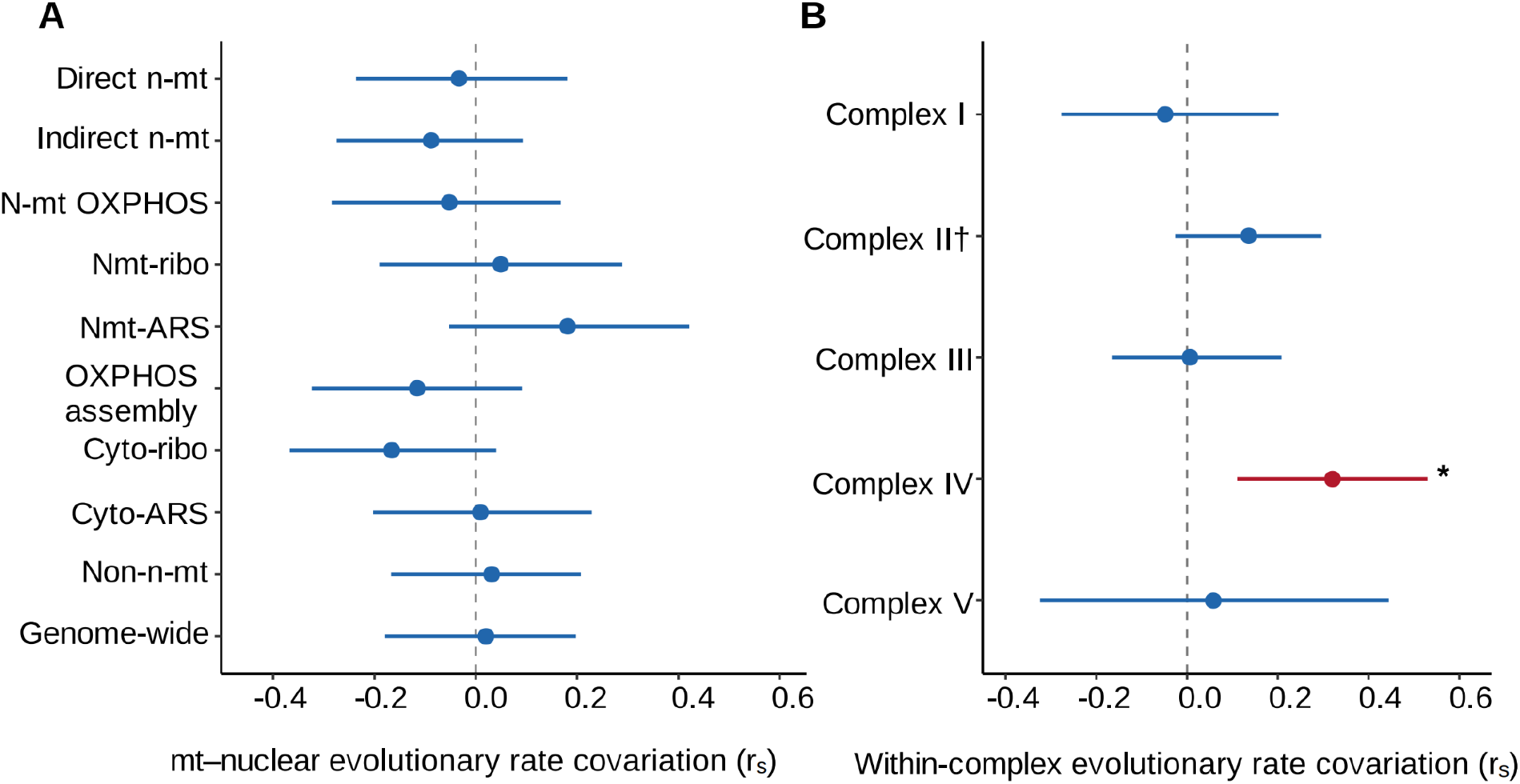
Evolutionary rate covariation between mitochondrial and nuclear gene sets. Evolutionary rate covariation (ERC) was quantified using Spearman correlations between branch-specific relative evolutionary rates. (A) Covariation between each nuclear gene set and the set of 13 mitochondrial protein-coding genes. For each branch, rates were averaged across genes within each set to obtain nuclear and mitochondrial composite-rate vectors, which were then correlated. (B) Within-complex covariation between mitochondrial- and nuclear-encoded OXPHOS subunits. The entirely nuclear-encoded Complex II was marked with a dagger (†). Points show mean Spearman correlation coefficients (r_s_) from 10,000 gene-bootstrap replicates, and horizontal bars show 95% bootstrap confidence intervals. Red indicates a positive ERC whose confidence interval excludes zero; blue indicates a confidence interval that includes zero. Asterisks indicate permutation support at *P* < 0.05 (*).

Non-n-mt genes, the nuclear genes with no known mitochondrial association, had a bootstrap mean correlation (r_s_) of 0.031 (95% CI = −0.167 to 0.208). The genome-wide nuclear reference set, which included all nuclear genes retained in the ERC analysis regardless of mitochondrial association, had a bootstrap mean correlation of 0.020 (95% CI = −0.179 to 0.198). Nmt-RPs and Nmt-ARSs had numerically more positive bootstrap mean correlations than their cytosolic counterparts (0.049 vs. −0.166 and 0.181 vs. 0.009, respectively), but neither mitochondrial translation-gene set showed a positive confidence interval excluding zero. Thus, none of the gene sets shown in Fig. 3A exhibited an overall positive evolutionary-rate correlation with mitochondrial rates whose bootstrap confidence interval excluded zero.

When nuclear OXPHOS genes were grouped by respiratory-chain complex, only Complex IV showed positive mitonuclear evolutionary-rate covariation (Fig. 3B). Evolutionary rates of the nuclear-encoded Complex IV subunits covaried with those of the mitochondrial-encoded COX1, COX2, and COX3 genes (observed r_s_ = 0.292; bootstrap mean r_s_ = 0.321, 95% CI = 0.110–0.530). The elevated covariation rate for Complex IV was not attributable to a single gene or phylogenetic branch. ERC remained positive after independently excluding each mitochondrial Complex IV gene (r_s_ = 0.299–0.333) or each nuclear Complex IV subunit (r_s_ = 0.209–0.392). Likewise, all leave-one-branch-out estimates were positive (r_s_ = 0.253–0.358). The observed association also exceeded expectations from both unrestricted branch-label permutations (*P* = 0.018) and permutations stratified between terminal and internal branches (Benjamini–Hochberg-adjusted *P* = 0.040). Together, these analyses identified Complex IV as the strongest candidate for mitonuclear evolutionary coupling.

### Population differentiation reveals recent mitonuclear concordance

Evolution of nuclear OXPHOS genes during stickleback’s colonization of freshwater habitats included both spatially restricted and widespread components. Population branch statistics (PBS) quantified population-specific allele-frequency differentiation relative to the corresponding marine population and a common outgroup (Fig. S6). Mean PBS across nuclear OXPHOS genes was significantly higher in British Columbia than in Alaska (median = 0.185 vs. 0.092; paired Wilcoxon signed-rank test, *P* = 1.13 × 10^-6^; Fig. 4A). Using the upper 5% of the regional PBS distributions as a descriptive criterion, *ndufs2*, *ndufb4* and *cox4i1* ranked in the upper 5% in both regions. *uqcrq* and *atp5pd* ranked in the upper 5% only in Alaska and occurred above the 1:1 line, whereas *cox6a1* and *hccsb* ranked in the upper 5% only in British Columbia and occurred below the 1:1 line. These rankings identify shared high-PBS candidates and regional asymmetry in the magnitude of differentiation, but top-5% membership alone was not interpreted as evidence of statistically significant region-specific effects.

**Figure 4.**
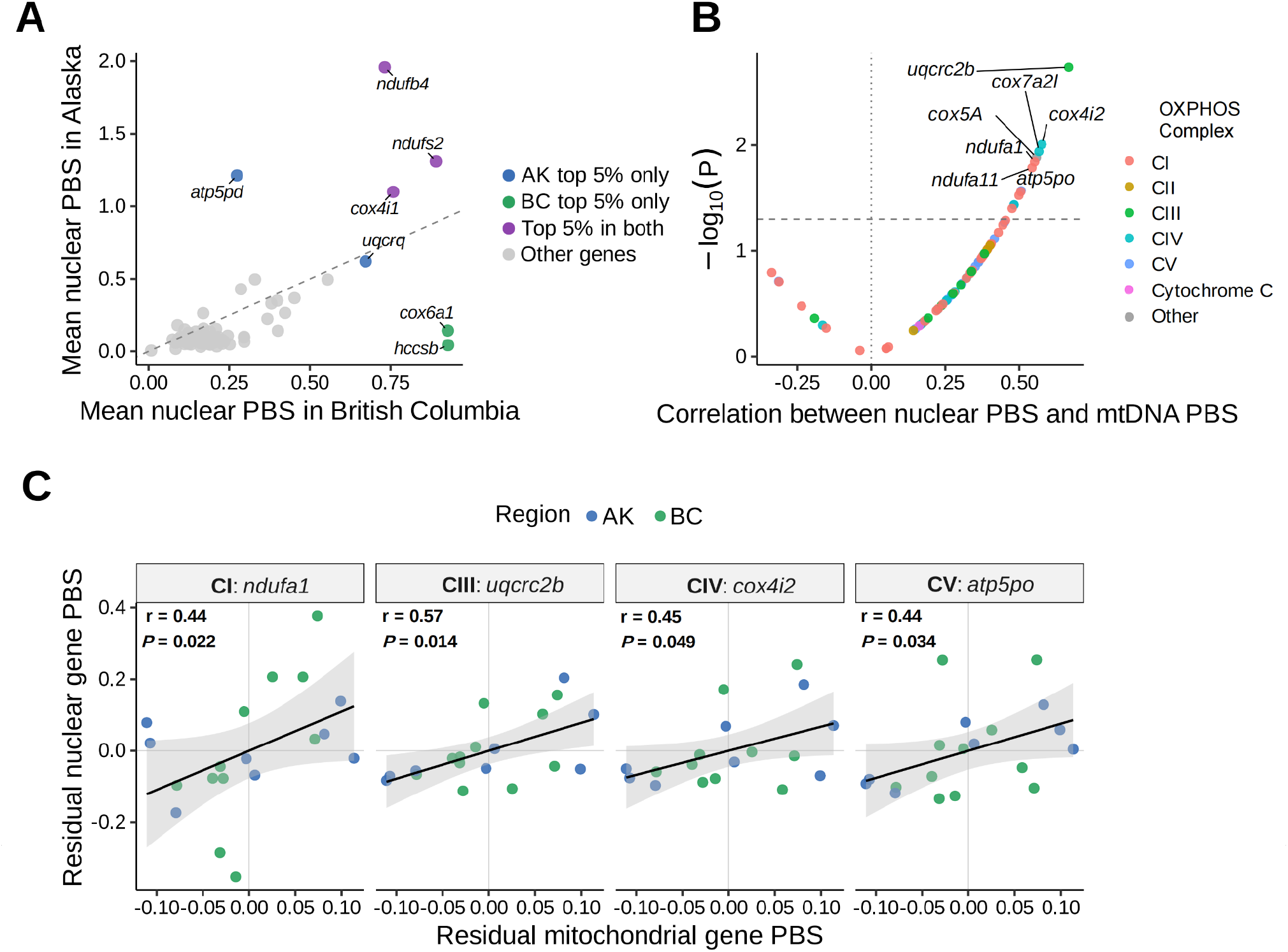
Regional differentiation and mitochondrial-associated variation in nuclear OXPHOS genes. (A) Mean nuclear population branch statistic (PBS) for each OXPHOS gene in Alaska and British Columbia. Each point represents one gene. Colored points indicate genes within the upper 5% of the PBS distribution in Alaska only, British Columbia only, or both regions. The dashed 1:1 line indicates equal mean PBS between regions; displacement from this line shows the magnitude and direction of regional differences. (B) Gene-wise Pearson correlations between nuclear and mitochondrial DNA PBS across freshwater populations. Points are colored by the OXPHOS complex. The vertical dotted line indicates zero correlation, and the horizontal dashed line indicates nominal significance (*P* = 0.05). (C) Relationships between nuclear and mitochondrial PBS after controlling for chromosome XXI PBS and geographic region. Genes with the strongest permutation-supported positive association within each OXPHOS complex (excluding Complex II) are shown. Points represent freshwater populations and are colored by region. Lines show fitted linear relationships, with shaded areas representing 95% confidence intervals. Residual correlations and *P* values from region-stratified permutation tests are reported within each panel.

Nuclear and mitochondrial PBS were broadly positively correlated across freshwater populations. Sixty-four of the 72 nuclear OXPHOS genes showed positive correlations, including 12 nominally significant associations (Pearson correlation, *P* < 0.05; Fig. 4B; Table S6). Among the 70 genes assigned to OXPHOS Complexes I–V, neither correlation strength nor the proportion of positively correlated genes differed significantly among complexes (gene-label permutation test, *P* = 0.994; Fisher’s exact test with Monte Carlo estimation, *P* = 0.930, respectively). Additionally, no individual complex was significantly enriched for positive correlations (one-sided Fisher’s exact tests, all BH-adjusted *q* = 0.918). However, similarly positive correlation distributions were observed for OXPHOS assembly factors, mitochondrial aminoacyl-tRNA synthetases and ribosomal proteins, and their cytosolic counterparts (Fig. S7). These widespread unadjusted associations therefore likely include effects of shared demographic history and genome-wide population structure. After controlling for geographic region and chromosome XXI PBS, and excluding Complex II which lacks mtDNA contributions, six nuclear OXPHOS genes retained positive associations with mitochondrial PBS in region-stratified permutation tests: *ndufa1*, *ndufa11*, *ndufa12*, *uqcrc2b*, *cox4i2*, and *atp5po* (*P* < 0.05; strongest association in each complex shown in Fig. 4C).

### Site-specific parallel allele-frequency changes in nuclear OXPHOS genes during freshwater colonization

We next tested whether changes in allele frequencies of nuclear OXPHOS genes showed parallel patterns across independent freshwater colonizations, and whether these responses varied with mitochondrial lineage. Although our PBS metrics identified genes with elevated freshwater-specific branch lengths, because the metric averages allele frequency shifts across all SNPs at a locus it may dilute evolutionary signatures at particular sites. We therefore identified the largest mean per-SNP allele-frequency change (max ΔAF) in each nuclear OXPHOS gene, averaged across populations, to test for parallel nuclear evolution during repeated freshwater colonization (Table S7). Per gene max ΔAF was strongly positively correlated between Alaska and British Columbia (Pearson’s r = 0.78, *P* = 6.2 × 10^-16^; Fig. 5A). Allele frequencies in 67 of 72 nuclear OXPHOS genes changed in the same direction in both regions, significantly exceeding the null expectation of 0.5 (two-sided binomial test, P = 6.39 × 10^-15^). Five genes showed large concordant shifts (|median ΔAF| ≥ 0.5) irrespective of geography, whereas another five reached this threshold in only one region (Fig. 5B). When considering the five genes with the most concordant and geographically widespread ΔAF patterns, there was a notable difference between populations likely to have colonized freshwater recently versus long-established populations. Specifically, the five recently colonized populations showed noticeably less divergence from marine ancestors. Overall, while we did observe some geographic structuring and variation in the magnitude of allele frequency shifts, we discovered five nuclear OXPHOS genes that display remarkably large and parallel evolution associated with freshwater habitats.

**Figure 5.**
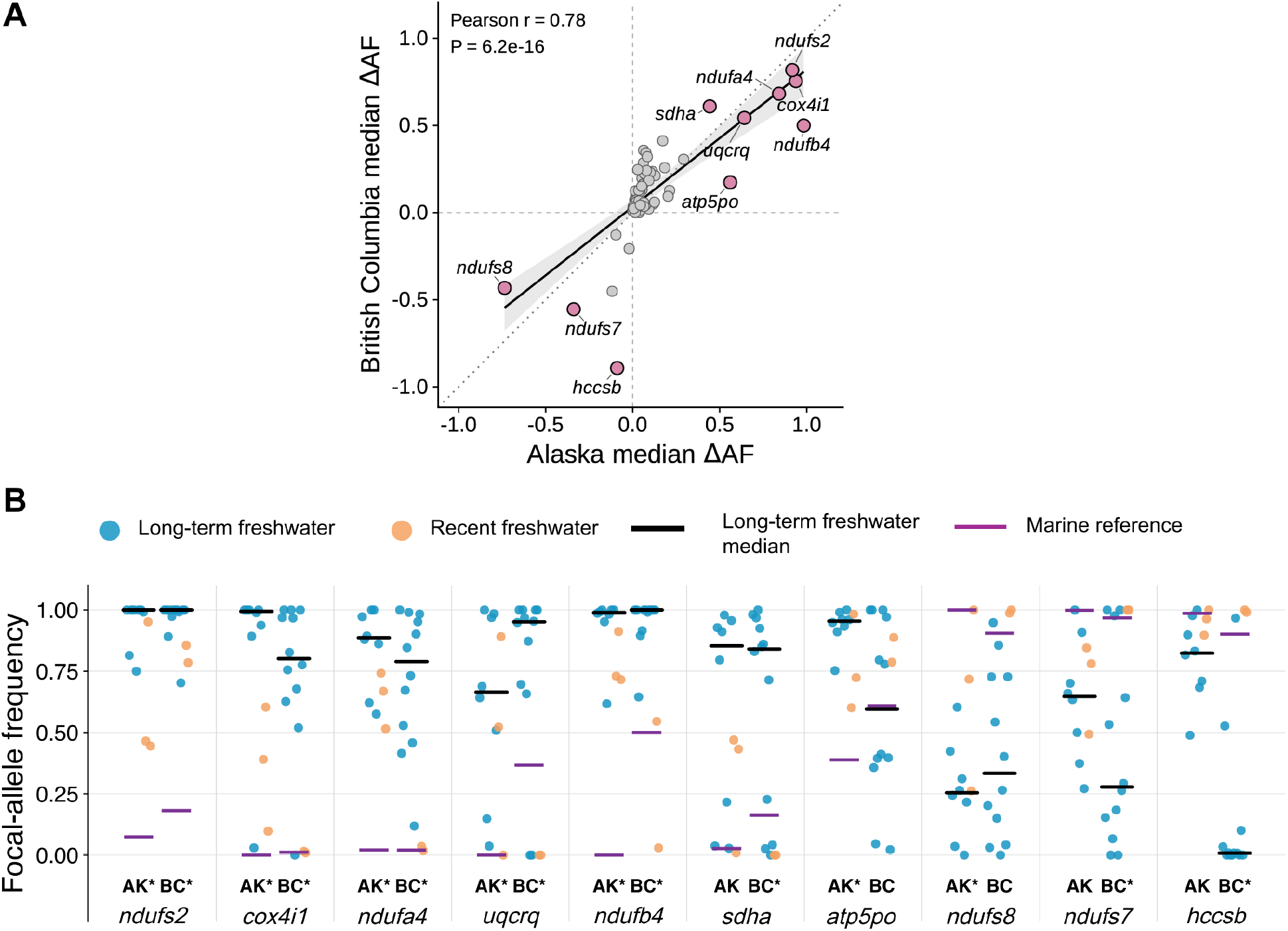
Parallel changes in nuclear OXPHOS allele frequencies across independently colonized freshwater populations in Alaska and British Columbia. (A) Gene-level allele-frequency shifts (ΔAF) in Alaska (AK) and British Columbia (BC), calculated as the regional median freshwater allele frequency minus the corresponding marine-reference frequency, including both long-term and recently colonized freshwater populations. Each point represents one of 72 nuclear OXPHOS genes, using the same representative SNP and focal allele in both regions. Pink points identify genes with |median ΔAF| ≥ 0.5 in at least one region. The solid line shows the linear regression with its shaded 95% confidence interval, and the dotted diagonal line indicates the 1:1 expectation. (B) Population-level focal-allele frequencies at representative SNPs for the ten candidate genes highlighted in panel A. Blue circles represent long-term freshwater populations, and orange circles represent the five recently colonized freshwater populations (SC, CH, LB, PACH, and FRED). Purple solid segments indicate focal-allele frequencies in the regional marine reference populations (RS for Alaska and SAY for British Columbia); black solid segments indicate median frequencies among long-term freshwater populations. Asterisks beside AK or BC indicate |median ΔAF| ≥ 0.5 in the corresponding region, using the all-population estimates shown in panel A.

### Geography and mitochondrial lineage explain distinct components of freshwater allele-frequency change

To further dissect patterns of evolutionary parallelism in the long-established freshwater populations we next tested whether variation among populations was associated with geographic location or mitochondrial lineage (excluding AMO). SNPs in strong linkage disequilibrium were pruned before analysis (Fig. S8), and geography and mitochondrial lineage were evaluated using separate SNP-level quasibinomial models.

Geographic associations were widespread among SNPs in the 72 nuclear OXPHOS genes (Fig. 6A; Table S8). Among all 41,062 OXPHOS SNPs tested across 19 established freshwater populations, 4,745 showed geographic associations at FDR (*q* < 0.05). Requiring a minimum fitted ΔAF of 0.20 retained 37 geography-associated candidate SNPs distributed across multiple nuclear OXPHOS genes. Mitochondrial-lineage associations were more restricted (Fig. 6B; Table S8). Specifically, 1,235 SNPs were associated with mitochondrial lineage at FDR (*q* < 0.05), but only three also showed a maximum fitted ΔAF of at least 0.20 among lineages. These SNPs occurred in *ndufa10*, *ndufab1b*, and *ndufs6*. All three genes encode Complex I subunits. No candidate SNP was shared between the geographic and mitochondrial-lineage analyses, although *ndufa10* contained different candidate SNPs in the two analyses.

**Figure 6.**
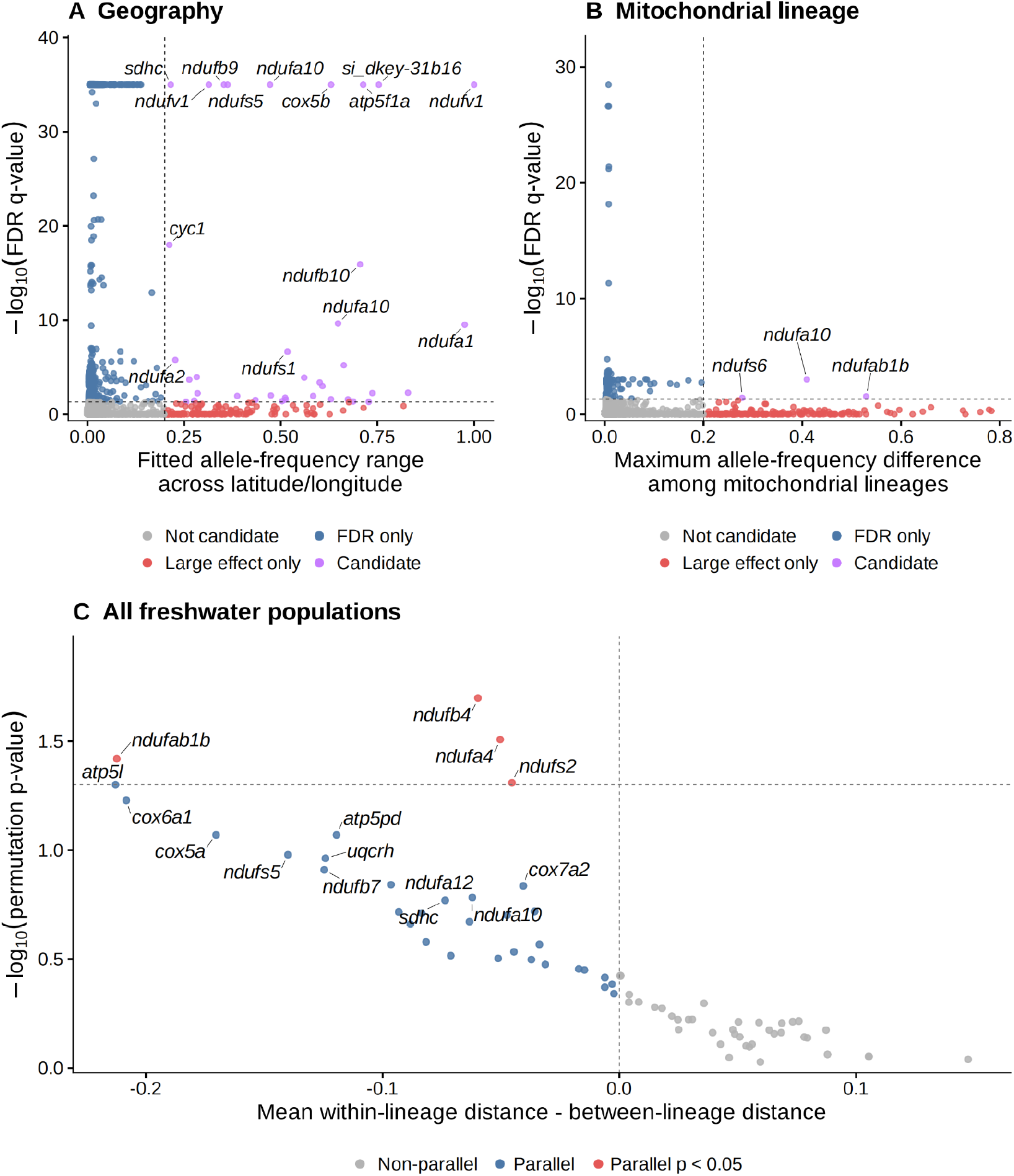
Geographic and mitochondrial-lineage structure of nuclear OXPHOS variation among freshwater populations. SNP-level quasibinomial generalized linear models tested associations between nuclear allele frequency and (A) geographic position or (B) mitochondrial lineage. Each point in (A) and (B) represents one SNP. The x-axis shows the maximum minus minimum fitted allele frequency across sampled geographic coordinates in (A) and the maximum fitted allele-frequency difference among mitochondrial lineages in (B). The y-axis shows −log_10_ of the FDR-adjusted *q*-value, capped at 35 for display. Purple points satisfy both *q* < 0.05 and an effect-size threshold of ≥0.20; blue points satisfy only the significance criterion, red points satisfy only the effect-size criterion, and gray points satisfy neither criterion. (C) Similarity of LD-pruned, signed ΔAF profiles among freshwater populations for each nuclear OXPHOS gene.

Gene-level ΔAF profiles provided additional evidence that mitochondrial lineage influenced a small subset of nuclear OXPHOS genes (Fig. 6C; Table S9). Four genes (*ndufb4*, *ndufa4*, *ndufab1b*, and *ndufs2*) showed greater permutation-supported similarity among populations sharing the same mitochondrial lineage than among populations from different lineages. Three of these genes (*ndufb4*, *ndufa4*, and *ndufs2*) also showed strong, concordant regional ΔAF in Alaska and British Columbia. Region-specific analyses identified more mitochondrial lineage-associated signals in British Columbia than in Alaska (Fig. S9).

Finally, we compared candidates identified by regional ΔAF, whole-gene PBS, and mitonuclear PBS association analyses. Only *atp5po* overlapped between the ten high-effect ΔAF genes and the six genes showing nominal positive associations between nuclear and mitochondrial PBS after controlling for geographic region and phylogenetic distance. Thus, genes showing large regional freshwater–marine allele-frequency differences were largely distinct from those whose differentiation covaried with mitochondrial PBS.

### Repeated freshwater colonization is associated with parallel amino-acid shifts in nuclear OXPHOS genes

To assess the potential functional consequences of parallel evolution in nuclear OXPHOS protein sequences, we identified 28 recurrent nonsynonymous substitutions across 72 nuclear OXPHOS genes, all of which passed read-depth, recurrence, and allele-frequency filters. Ten substitutions in five genes showed concordant shifts with |median ΔAF| ≥ 0.2 and permutation *P* < 0.05 in both Alaska and British Columbia. Applying |median ΔAF| ≥ 0.5 in both regions retained eight substitutions in four genes (*cox4i1*, *ndufs2*, *ndufs8*, and *sdha*; Table S10), all of which were also highlighted as candidates in Fig. 5. Despite their large parallel frequency shifts, most of the 28 substitutions were classified as tolerated by SIFT4G. Five were predicted to be deleterious (SDHA D597N and D535V, UQCRC2B L8P and A2V, and DMAC2L L10V), and three had BLOSUM62 scores ≤ −2. Among the eight substitutions with |median ΔAF| ≥ 0.5 in both regions, only SDHA D597N was predicted to be deleterious, and none had a BLOSUM62 score ≤ −2.

Among the previously identified candidate genes, *cox4i1* was the only locus that combined cross-region PBS outlier status, large parallel allele-frequency shifts, and three nonsynonymous substitutions with |median ΔAF| ≥ 0.5 in both Alaska and British Columbia. These variants corresponded to A58V, F124Y, and F133L relative to the marine-associated amino-acid states. All three were classified as tolerated by SIFT4G (scores of 1.0, 1.0, and 0.5, respectively) and had BLOSUM62 scores of 0, 3, and 0, respectively (Table S10). *cox4i1* was also among the most extreme upper-tail outliers for both d_N_ and d_N_/d_S_ among nu OXPHOS genes. In addition, COX4I1 is a component of Complex IV, the only OXPHOS complex showing positive evolutionary-rate covariation with a bootstrap confidence interval excluding zero (observed Spearman’s r_s_ = 0.292; bootstrap mean = 0.321, 95% CI = 0.110–0.530).

Given these signals, we asked whether the three substitutions are likely to affect protein function by examining cross-species conservation and predicted structural interactions with mitochondrially encoded Complex IV subunits. The freshwater-associated V58, Y124, and L133 states increased in frequency in both Alaska and British Columbia, although their frequencies varied among freshwater populations (Fig. 7A,B). AlphaFold-Multimer models containing COX4I1, MT-CO1, MT-CO2, and MT-CO3 produced similar overall architectures for the representative marine- and freshwater-associated COX4I1 sequences (Fig. 7C,D). Models were high-confidence (Complex ipTM 0.88–0.90 for both sequence states across five models each), and confidence did not differ appreciably between states. Across five matched model pairs, residue 58 remained distant from the mitochondrially encoded subunits (approximately 21–23 Å), whereas residue 124 occurred near MT-CO1 (approximately 7.6 Å) but differed little between states. Residue 133 formed the closest predicted mitonuclear contact. The marine-associated F133 was approximately 3.2–3.3 Å from MT-CO1, compared with 3.7–3.9 Å for the freshwater-associated L133 residue. Aligning the freshwater and marine COX4I1 amino acid sequences against eight other vertebrates revealed that residue 58 was highly variable across species, while the other two positions were moderately conserved (Fig. 7E). Specifically, positions 124 and 133 each had two amino acid states with similar biochemical and structural properties, but the freshwater stickleback states were observed across multiple divergent species.

**Figure 7.**
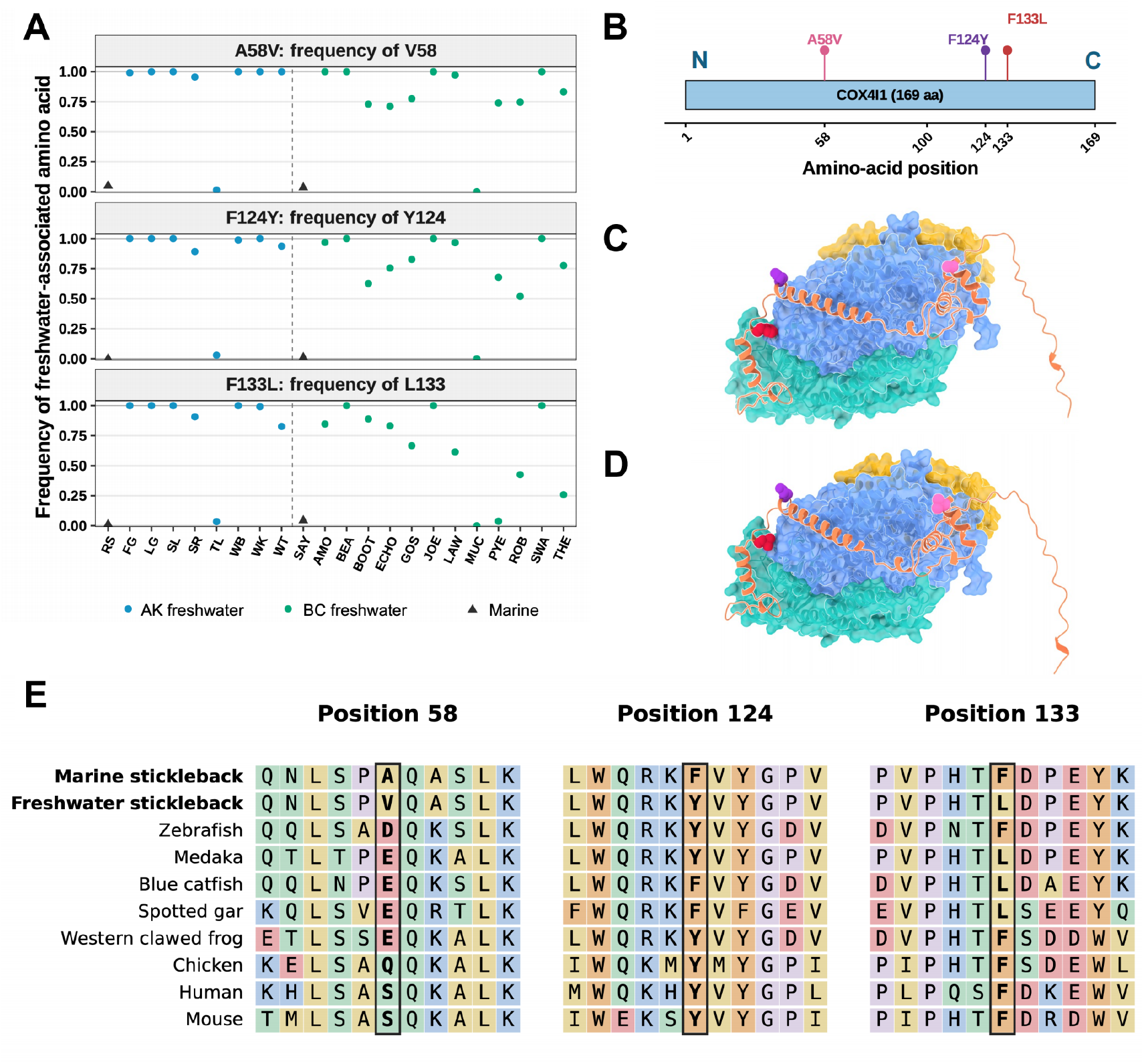
Freshwater-associated COX4I1 amino-acid variants: population frequencies, cross-species conservation, and predicted structural positions. (A) Frequencies of the freshwater-associated V58, Y124, and L133 states across two marine reference populations and 20 freshwater populations. Populations are ordered by region, with the vertical dashed line separating Alaska from British Columbia. Frequencies were reconstructed from strand-aware Pool-seq allele counts using the canonical *cox4i1* coding sequence. (B) Positions of the three focal substitutions along the 169-amino-acid COX4I1 protein with color coding to match their locations in 3D structural predictions. N and C indicate the N- and C- termini, respectively. (C, D) Rank-1 AlphaFold-Multimer models of the marine-associated (C) and freshwater-associated (D) COX4I1 sequence variants with MT-CO1, MT-CO2, and MT-CO3. COX4I1 is shown as an orange ribbon; MT-CO1, MT-CO2, and MT-CO3 are shown as blue, teal, and gold surfaces, respectively. Both models show similar overall architectures. (E) Local amino-acid alignments surrounding positions 58, 124, and 133 in the marine-associated and freshwater-associated stickleback sequences and eight other vertebrate species. Residues are colored by physicochemical class, and black boxes mark the focal sites. Position numbers refer to the ungapped stickleback sequence.

## Discussion

Mitochondrial and nuclear genomes are expected to coevolve to maintain essential cellular functions (Sloan et al. 2018) and potentially mediate adaptive responses to environmental shifts (Hill 2015). The repeated colonization of freshwater habitats by threespine stickleback potentially provides an opportunity to examine both of these processes across populations with diverse mitochondrial backgrounds. We first documented extensive variation in mitochondrial and nuclear genes involved in mitochondrial function across marine and freshwater populations, establishing the utility of this system for examining molecular evolution during repeated environmental adaptation. We then identified evidence of mitonuclear coupling, elevated nonsynonymous divergence and diversity in mitochondrial translation genes, and pronounced parallel allele-frequency shifts in nuclear OXPHOS genes associated with freshwater colonization.

The weak geographic clustering and extensive divergence of mitochondrial population-consensus sequences (Fig. 1A) suggest that freshwater colonization involved differential sorting of pre-existing mitochondrial variation. Given that most freshwater populations in these formerly glaciated regions were established within the past approximately 10,000–15,000 years (Deagle et al. 2013; McGee and Wainwright 2013), much of this divergence likely predates freshwater colonization rather than having arisen entirely through subsequent de novo mutation. This interpretation is consistent with evidence for persistence in Pleistocene refugia (Ortí et al. 1994), postglacial dispersal (Johnson and Taylor 2004) and admixture among mitochondrial lineages (Liu et al. 2016). Differential sorting of this ancestral variation could reflect founder effects and genetic drift, but could also arise through localized selection acting on a diverse founding pool. Watershed membership did not significantly predict mitochondrial cluster membership after controlling for region, nor did it improve prediction relative to region alone. Amor lake (AMO) provides a complementary observation: although excluded from the cluster-based analysis because of its exceptional divergence, it shares the Amor River watershed with Robert lake (ROB), which belongs to the mitochondrial cluster containing the marine reference populations. Their contrasting mitochondrial backgrounds challenge a simple model in which a shared founder event during upriver colonization imposed the same mitochondrial lineage throughout the watershed. Separate colonization events and subsequent demographic processes could also produce this contrast, but localized selection on standing mitochondrial variation merits consideration alongside these explanations.

The population-consensus sequences of the two marine reference populations occurred near the center of the mitochondrial network, whereas freshwater populations were distributed across several differentiated groups. This pattern is compatible with differential retention or selection of ancestral mitochondrial variation during freshwater colonization. However, the small number of marine reference populations and the limitations of Pool-seq prevent us from resolving individual haplotypes, estimating their frequencies, or determining whether the observed differentiation exceeds expectations under colonization bottlenecks and genetic drift. Selection on standing mitochondrial variation therefore remains a plausible explanation, but cannot be established from these patterns alone. AMO additionally carried an exceptionally divergent mitochondrial background and formed a distinct branch relative to the other northeastern Pacific populations in our phylogenetic analysis (Fig. 1B). Previous studies have documented deeply divergent mitochondrial lineages in stickleback populations from Alaska and British Columbia (Ortí et al. 1994; Deagle et al. 1996; Thompson et al. 1997; Johnson and Taylor 2004). Mitochondrial introgression from Japan Sea stickleback has been proposed as an explanation for these divergent lineages (Beck et al. 2022), and future analyses of the AMO haplotype could test whether it conforms with this model.

Against this heterogeneous mtDNA background, patterns of molecular evolution in nuclear genes differed markedly among OXPHOS and mitochondrial-translation gene classes (Fig. 2). Mitochondrial translation genes showed a clearer contrast with their cytosolic counterparts in both divergence and within-population diversity (Figs. 2B,C and S2B,C). Both Nmt-RPs and Nmt-ARSs exhibited higher nonsynonymous divergence and diversity without corresponding increases in synonymous measures. The Nmt-ARS pattern was particularly pronounced: median d_N_ and d_N_/d_S_ exceeded both the corresponding cyto-ARS values and the genome-wide nuclear medians, indicating that the contrast was not driven solely by unusually slow evolution of cyto-ARSs. Similar patterns of accelerated evolution of nuclear-encoded mitochondrial translation proteins have been reported across diverse taxa. In the copepod *T. californicus*, the Nmt-RPs accumulated more nonsynonymous substitutions than cytosolic RPs (Barreto and Burton 2013). The authors interpreted this pattern as consistent with compensatory responses to rapidly evolving mitochondrial rRNAs. A subsequent genome-wide analysis of eight *T. californicus* populations found elevated evolutionary rates and signatures of positive selection in nuclear genes predicted to interact with mtDNA or its encoded proteins and RNAs across multiple pathways, supporting a broader role for compensatory mitonuclear evolution (Barreto et al. 2018). Higher d_N_ and d_N_/d_S_ in Nmt-ARSs than in cytosolic ARSs have also been reported in mammals, birds, and *Drosophila* (Adrion et al. 2016), with related patterns found in nonbilaterian animals (Pett and Lavrov 2015) and organelle-targeted ribosomal proteins of *Silene* plants (Sloan et al. 2014). Thus, the elevated nonsynonymous evolution of stickleback Nmt-ARSs, together with their functional interactions with mitochondrially encoded tRNAs, is consistent with compensatory selection to maintain mitonuclear function.

In contrast to Zhang and Broughton (2013), who reported lower d_N_ in nuclear core than noncore OXPHOS subunits across vertebrates, within stickleback we detected higher d_N_ and d_N_/d_S_ in nuclear core subunits (Fig. S4A). Because these core subunits interact directly with mtDNA-encoded proteins to generate the catalytic capacities of OXPHOS complexes, their elevated rate of divergence indicates another potential instance of compensatory mitonuclear evolution. Notably, we did not observe elevated rates of divergence when comparing direct n-mt with indirect n-mt or non-n-mt (Fig. S5A). Similarly, Kuster et al. (2026) found mitonuclear ancestry did not differ consistently among these broad gene classes in hybridizing swordtails, despite known large-effect mitonuclear incompatibilities. Such results suggest that mitonuclear selection may be concentrated in particular genes or functional modules rather than distributed uniformly across broad mitochondrial gene classes.

Evolutionary-rate covariation analyses provided more localized evidence of evolutionary coupling between mitochondrial and nuclear genes in particular functional modules (Fig. 3). Nmt-ARSs and Nmt-RPs tended to show more positive rate covariation than their cytosolic counterparts, but these differences were weak, and direct n-mt genes did not collectively exceed indirect n-mt genes or the broader nuclear background (Fig. 3A). Rate covariation was more clearly resolved at the level of individual OXPHOS complexes. Complex IV showed the strongest within-complex signal, which remained positive after removing individual genes or phylogenetic branches (Fig. 3B). Its mtDNA-encoded COX1–3 subunits form the catalytic core and function closely with nuclear-encoded subunits, making Complex IV a plausible target of mitonuclear coevolution (Osada and Akashi 2012). The entirely nuclear-encoded Complex II showed the second-highest, although modest, covariation with mitochondrial evolutionary rates. Because Complex II contains no mtDNA-encoded subunits, this pattern cannot reflect within-complex mitonuclear coevolution. Instead, it may reflect coordinated selection on respiratory metabolism or the broader roles of Complex II in metabolic regulation, succinate signaling, inflammation, and cell fate (Iverson et al. 2023). Inflammation also varies heritably among stickleback populations (Weber et al. 2017; Weber et al. 2022), though its connections to Complex II are unknown. The restriction of clear within-complex rate covariation to Complex IV contrasts with the broader mitonuclear patterns reported across deeply diverged insects and mammals (Yan et al. 2019; Weaver et al. 2022). This difference may reflect evolutionary timescale, because the recent postglacial divergence of stickleback populations provides less time for coordinated substitutions to accumulate than the deeper interspecific divergences examined in those studies. However, many freshwater-adaptive alleles in threespine stickleback derive from ancient standing genetic variation (Jones et al. 2012; Nelson and Cresko 2018). Thus, future research estimating the ages of freshwater-associated OXPHOS alleles could determine whether the limited mitonuclear coupling reflects their relatively short history in freshwater or a lack of functional interdependence.

Whereas evolutionary-rate covariation analyses measured covariation in substitution rates across phylogenetic branches, population branch statistic metrics allowed us to test whether nuclear and mitochondrial loci show concordant allele-frequency differentiation among recently diverged populations. Nuclear and mitochondrial PBS were predominantly positively correlated (Fig. 4B), but similar patterns among nonmitochondrial comparison genes indicated that much of this concordance reflected shared demographic and geographic history (Sloan et al. 2015; Zaidi and Makova 2019; Fig. S7). Nevertheless, several nuclear OXPHOS associations remained after accounting for region and genome-wide differentiation (Fig. 4C). The strongest demographic corrected association within Complex IV occurred at *cox4i2*, and several other nuclear Complex IV genes showed elevated freshwater PBS or positive associations with mitochondrial PBS. Notably, phylogenetic rate covariation and recent population differentiation both implicated Complex IV, although they identified different individual genes. This incomplete overlap is expected because the two analyses capture distinct evolutionary timescales: rate covariation summarizes substitutions across phylogenetic branches, whereas PBS measures recent allele-frequency differentiation among populations. We therefore next tested whether recent differentiation involved parallel changes at the same SNPs across independent freshwater colonizations.

Nuclear OXPHOS allele-frequency shifts were strongly correlated between Alaska and British Columbia (Fig. 5A). Genes showing large freshwater–marine differences in one region generally exhibited similarly directed differences in the other, consistent with repeated responses to freshwater selection. Stickleback adaptation frequently draws on standing nuclear genetic variation maintained at low frequencies in marine populations, enabling rapid and geographically replicated evolutionary responses (Colosimo et al. 2005; Jones et al. 2012; Lescak et al. 2015; Nelson and Cresko 2018). Many OXPHOS variants followed this pattern: they were rare in the regional marine populations but increased dramatically across numerous freshwater populations (Fig. 5B). None of the examined OXPHOS genes reside in the three chromosomal inversions associated with freshwater adaptation across many stickleback populations (Jones et al. 2012; Fang et al. 2020; Roberts Kingman et al. 2021). However, a recent analysis, using largely distinct populations from those examined in this paper, identified a suite of genomic regions showing recurrent marine–freshwater differentiation in the Northeast Pacific region (Roberts Kingman et al. 2021). All ten genes in Fig. 5B overlapped with these differentiated regions. Moreover, three OXPHOS genes (*uqcrq*, *atp5po*, and *ndufs8*) are in close proximity (0.95–3.59 Mb) to the gene *eda* on chromosome four, which is perhaps the most studied stickleback locus with respect to freshwater adaptation (Colosimo et al. 2005; Barrett et al. 2008; Jones et al. 2012). While selection on the freshwater allele of *eda* may drive some of differentiation at nearby OXPHOS genes, an alternative explanation is that concerted selection on each locus, combined with recombination, creates a large freshwater haplotype that facilitates the strong and rapid adaptation observed in nature (Roberts Kingman et al. 2021). This latter hypothesis warrants additional attention in the light of our new evidence for genome-wide selection on OXPHOS genes.

While many OXPHOS loci displayed largely parallel shifts, some deviated from this general pattern. In contrast to the large marine-freshwater differentiation of OXPHOS genes in long-term freshwater populations, recently colonized freshwater populations often exhibited intermediate or a lack of divergence from the marine state (Fig. 5B). Populations with intermediate frequencies may represent early or ongoing stages of freshwater adaptation, and those lacking marine divergence may inhabit environments that impose more marine-like selective pressures. However, limited information on population ages across our dataset prevented us from further investigating these hypotheses. There were also a handful of genes that displayed high levels of divergence in either Alaska or British Columbia, not in both regions (Figs. 4, 5, S9). Future work could examine whether this disparity results from environmental differences between the regions, or potentially from the loss of genetic variation due to bottlenecks as stickleback migrated out of their ancestral range (Fang et al. 2020; Roberts Kingman et al. 2021).

Apart from assessing (non) parallelism in OXPHOS evolutionary trends, we also compared the relative influences of mtDNA lineage and geography. Geographic associations were more widespread than mitochondrial-lineage associations among nuclear OXPHOS SNPs (Fig. 6). The limited overlap between geography- and mitochondrial lineage-associated candidates further suggests that these factors were associated with largely distinct components of nuclear OXPHOS variation. Mitochondrial lineage associations were concentrated in a smaller subset of nuclear OXPHOS loci, highlighting candidates for potential mitonuclear coadaptation, through which the effects can depend on mitochondrial background (Rand et al. 2004; Dowling et al. 2008; Burton and Barreto 2012). All three lineage-associated candidate SNPs belong to Complex I. Complex I integrates mitochondrial and nuclear gene products and couples NADH oxidation and ubiquinone reduction to proton translocation (Kampjut and Sazanov 2020), while also contributing to reactive oxygen species production and redox signaling (Murphy 2009). These functional properties may provide a mechanistic lens for examining how environmental conditions and mitochondrial background shape variation in Complex I.

Finally, given the substantial nucleotide- and coding-level differences that we observed in OXPHOS genes, we focused on a single gene that stood out across comparison and tested whether amino acid shifts were predicted to alter protein function. Our focal gene, nuclear-encoded isoform 1 of cytochrome *c* oxidase subunit 4 (*cox4i1*), provides a plausible link between freshwater adaptation and mitonuclear evolution because its protein product contacts the mitochondrially encoded catalytic core of Complex IV (Sinkler et al. 2017). Moreover, COX4I1 contributes to the regulation of Complex IV activity, including responses to cellular energy status through adenine-nucleotide binding (Arnold and Kadenbach 1997; Acin-Perez et al. 2011). Our computational predictions found that freshwater-associated amino acid changes had little effect on the protein structure of COX4I1 or its interactions within Complex IV, nor were these residues highly conserved across vertebrates (Fig. 7). However, these results cannot exclude effects on protein stability, dynamics, or allosteric regulation. Variation in COX4I1 may influence respiratory function through changes in subunit interactions or regulation, even in the absence of pronounced predicted structural changes. For example, we did observe a small difference in predicted intersubunit distance at residue 133. Alternatively, the increased frequency of freshwater-associated coding variants could reflect selection on linked noncoding variants that affect gene expression. In this context, we note that the variant nearest the 5’ end of the *cox4i1* showed the greatest freshwater divergence, particularly in British Columbia populations. Future functional experiments are necessary to confirm and distinguish between these alternative possibilities.

## Conclusion

Together, our results reveal that large, parallel allele-frequency shifts in nuclear OXPHOS genes evolved across independently colonized freshwater stickleback populations. This is consistent with common ecological selection on mitochondrial function, although linked selection cannot be excluded. Mitochondrial DNA lineage was associated with a smaller, locus-specific component of nuclear OXPHOS variation, concentrated in Complex I genes. Elevated nonsynonymous divergence and diversity in mitochondrial translation genes suggest that selective constraints differ among functional modules, whereas rate covariation within Complex IV provides more localized evidence of mitonuclear coupling. Genes with the largest freshwater-associated shifts overlapped little with those whose differentiation covaried with mitochondrial DNA, suggesting that selection associated with freshwater colonization and mitonuclear coupling may act largely on different loci. These results highlight the importance of both ecological transitions and mitochondrial background in shaping nuclear OXPHOS evolution, and show that threespine stickleback is a useful system for studying how these factors jointly influence adaptation.

## Materials and Methods

### Study populations and sequencing data

We analyzed previously published pooled whole-genome sequencing data from 27 threespine stickleback (*Gasterosteus aculeatus*) populations sampled across Alaska and British Columbia (Weber et al. 2022). The dataset comprised eight established freshwater populations from Alaska, 12 established freshwater populations from British Columbia, two regional anadromous marine populations, and five recently colonized young freshwater populations (PACH, FRED, SC, LB, and CH) known or inferred to have been founded or recolonized by anadromous ancestors within the past several decades (Lescak et al. 2015; Aguirre et al. 2022). RS and SAY were used as the marine references for Alaska and British Columbia, respectively. Geographic coordinates, watershed assignments, habitat classifications, pool sizes, mitochondrial-lineage assignments, and inclusion in individual analyses are reported in Table S1. All 27 populations were included in analyses of mitochondrial phylogeny and nucleotide diversity. AMO was excluded from mitochondrial ordination and analyses based on discrete mitochondrial clusters because its exceptional mitochondrial divergence exerted disproportionate influence on the mitochondrial distance structure.

### Reference genomes and sequence processing

Mitochondrial analyses used the threespine stickleback mitochondrial reference genome MH205729.1 (Jiang et al. 2018). Nuclear analyses used the GAculeatus_UGA_version5 threespine stickleback assembly and its corresponding GFF3 annotation (Nath et al. 2021). Raw paired-end reads were processed with BBDuk in BBTools v38.90 using ktrim=rl, trimq=20, minlength=25, and ftl=10. Read quality was evaluated with FastQC v0.12.1 (Andrews 2010). Reads were aligned separately to the mitochondrial and nuclear references using Bowtie2 v2.5.4 (Langmead and Salzberg 2012). Both alignments applied no-mixed, no-discordant, and a maximum fragment length of 2,000 bp; mitochondrial alignments additionally used a very-sensitive-local option. Alignments with mapping quality <20 were removed, and BAM files were sorted and indexed with SAMtools (Li et al. 2009). Duplicate reads were removed from nuclear alignments using Picard but were retained in mitochondrial alignments. Mitochondrial variants were independently called using GATK HaplotypeCaller v4.6.1.0 (Poplin et al. 2017), with ploidy set to the number of individuals in each pool, and BCFtools v1.17 (Li 2011) in haploid mode. Per-population GATK and BCFtools calls were normalized, and records with genotypes containing the alternative allele were retained before intersection. Biallelic SNPs with matching reference and alternative alleles in both callsets were retained, requiring BCFtools read depth of 10–5,000 and variant quality ≥30. Potential NUMT interference was assessed by remapping reads to a combined nuclear and mitochondrial reference and comparing the resulting mitochondrial consensus sequences and lineage assignments with the original results.

### Mitochondrial consensus sequences and population relationships

Population-level mitochondrial consensus sequences were generated using BCFtools by applying filtered haploid SNP calls to the mitochondrial reference. Site-level allelic depths were calculated using reads with mapping quality ≥20 and bases with quality ≥13. Positions with usable depth <10 or >5,000, or without a nucleotide accounting for at least 80% of the allelic depth, were masked as N. Additional masks covered rejected alternative-allele candidates, predominant alternative alleles lacking retained SNP support, and positions where retained SNP calls disagreed with the predominant pileup base. Candidate indels and their 5-bp flanking regions were also masked. Only SNP substitutions were incorporated, preserving reference coordinates. Consensus sequences were aligned using MAFFT v7.525 (Katoh and Standley 2013). These sequences summarize population-level mitochondrial variation and do not resolve individual haplotypes or within-population haplotype frequencies. The control region was excluded from phylogenetic analyses because of its high indel content and alignment uncertainty.

A maximum-likelihood phylogeny was reconstructed from the control-region-excluded mitochondrial alignment using IQ-TREE v2.4.0 under the GTR+G substitution model (Nguyen et al. 2015). Branch support was evaluated using 1,000 ultrafast bootstrap replicates and SH-aLRT tests. The mitochondrial chrM sequence extracted from the Japan Sea stickleback (*Gasterosteus nipponicus*) assembly JS_male_hap1.v20240517 was included as the outgroup, and the tree was rooted on this sequence (Yamasaki et al. 2025). Internal branch support values were displayed when both SH-aLRT ≥80% and ultrafast bootstrap ≥95%. A median-joining network was constructed from the control-region-excluded population alignment with PopART v1.7 using default settings (Bandelt et al. 1999; Leigh and Bryant 2015). Indels and ambiguous positions were treated as missing.

Pairwise patristic distances among the study populations were extracted from the mitochondrial maximum-likelihood tree using ape v5.8-1. Principal coordinate analysis was performed using cmdscale in R after excluding AMO because its exceptional divergence strongly influenced the ordination. The Japan Sea outgroup was excluded from ordination and clustering. Mitochondrial population groupings were evaluated using *k*-means clustering, and four clusters (C1–C4) were used for downstream nuclear analyses. Population assignments are reported in Table S1.

We tested whether watershed grouping predicted mitochondrial cluster membership among the 24 freshwater populations with cluster assignments, including the five recently colonized populations. AMO and the two marine reference populations were excluded. Watershed assignments were obtained from Table S1. Association was evaluated using the sum of within-region Pearson chi-square statistics, with significance assessed by 9,999 permutations of cluster labels within Alaska and British Columbia, thereby preserving regional cluster frequencies. Predictive performance was evaluated by leave-one-population-out cross-validation. Each held-out population was assigned the most frequent cluster among the remaining populations from the same region and watershed. Prediction was evaluated for the 19 populations with at least one such training population; populations from singleton watersheds were not scored. Ties among the most frequent clusters received fractional credit. Watershed-based accuracy was compared with a region-only majority-class predictor evaluated on the same held-out populations, and the accuracy gain was tested using the same within-region permutation procedure.

### Mitochondrial differentiation and diversity

Mitochondrial allele counts were generated jointly across populations using SAMtools and converted to synchronized format using PoPoolation2 (Kofler et al. 2011). Minimum mapping and base qualities were both set to 30, and the maximum read depth was set to 5,000 per position per population during pileup generation. Pairwise *F*_ST_ was estimated using the unbiased Nei estimator implemented in grenedalf v0.6.3, with a minimum allele count of two and minimum depth of four (Czech et al. 2024). Estimates were obtained for the complete mitochondrial genome and separately for annotated protein-coding genes, rRNAs, tRNAs, and the control region using BED-defined intervals. Nucleotide diversity (π) was calculated in 500-bp sliding windows with a step size of 100 bp and separately for annotated mitochondrial regions. Population-specific pool sizes were supplied to grenedalf for finite-sample correction.

### Nuclear population structure

Nuclear population structure was summarized using the previously reconstructed chromosome XXI population tree based on *F*_ST_ values (Weber et al. 2022). Pairwise patristic distances extracted from this tree were subjected to principal coordinate analysis. The first two axes, treePC1 and treePC2, explained 35.12% and 14.32% of the variation, respectively, and were included as covariates in mitochondrial-cluster association models. Chromosome XXI PBS was used as an empirical background measure of nuclear population differentiation in covariate-adjusted PBS analyses, without assuming that chromosome XXI was entirely neutral.

### Functional gene sets and orthology assignment

We retained the genome-wide nuclear gene set and corresponding functional categories (direct n-mt, indirect n-mt, and non-n-mt) defined by Kuster et al. (2026). Under this classification, direct n-mt genes encode products with documented physical or molecular interactions with mitochondrially encoded proteins or RNAs; indirect n-mt genes encode mitochondrially localized products without known direct interactions with mitochondrially encoded components; and non-n-mt genes lack annotated mitochondrial functions.

We also curated nuclear gene sets representing OXPHOS structural subunits, OXPHOS assembly factors, mitochondrial aminoacyl-tRNA synthetases (Nmt-ARSs), mitochondrial ribosomal proteins (Nmt-RPs), and functionally analogous cytosolic aminoacyl-tRNA synthetases (cyto-ARSs) and ribosomal proteins (cyto-RPs). MitoCarta3.0 provided the principal reference for mitochondrial localization and function (Rath et al. 2021); cyto-ARSs and cyto-RPs were obtained from MSigDB and HGNC, respectively (Liberzon et al. 2011; Tweedie et al. 2021).

Human gene names and their teleost-specific aliases were first matched to the stickleback annotation. Unresolved loci were identified by querying human proteins against the predicted stickleback proteome with DIAMOND blastp using the more-sensitive option (Buchfink et al. 2015). Candidate orthologs were evaluated using reciprocal support, alignment coverage, protein length, gene-model integrity, and consistency with expected function.

For curated gene sets initially defined using human gene symbols, a representative stickleback ortholog was selected using the criteria above. Additional paralogous loci remained independent genes in genome-wide analyses. Loci were tracked by stable Ensembl gene identifiers and genomic coordinates rather than by potentially ambiguous gene symbols. Gene identities and paralog relationships were cross-checked against the GAculeatus_UGA_version5 GFF3 annotation and OMA orthology and paralogy records accessed through Bgee. Alternative transcripts sharing the same Ensembl gene identifier were treated as isoforms rather than separate paralogs.

Nuclear OXPHOS structural subunits were classified as core or noncore following Zhang and Broughton (2013). Core subunits have homologs in bacterial respiratory complexes and generally contribute directly to catalysis, electron transport, or proton translocation. Noncore subunits are later-derived accessory components associated with assembly, stability, or regulation. Assembly factors were retained as a separate class.

Mean CDS coverage was estimated with mosdepth v0.3.1 and normalized to median autosomal coverage (Pedersen and Quinlan 2018). Genes with normalized coverage outside 0.7–1.3 were inspected as possible copy-number, mapping, or orthology anomalies. Gene membership and numbers retained after each analysis-specific filter are reported in Table S2.

### Codon alignments

For each nuclear gene, a representative transcript was obtained from the predefined gene annotation. Population-specific consensus sequences were generated by incorporating high-confidence biallelic SNPs into the nuclear reference genome using BCFtools. Retained alternative SNPs required variant quality ≥30, depth ≥10, and an alternative-allele fraction ≥0.80. Positions with depth <10 were masked, together with positions affected by variants that failed these filters. Coding sequences were extracted from the masked consensus genomes using the same transcript coordinates and strand orientation across populations. Nuclear codon alignments were constructed by matching homologous CDS positions across all 27 populations. Sequences were required to have equal lengths divisible by three. Codons containing ambiguous bases or stop codons were masked, and the primary divergence analyses retained only codon columns that were complete and unambiguous in every population. Gene alignments were required to retain at least 100 complete codons and at least 50% of the original codon positions. The 13 mitochondrial protein-coding genes were extracted from the population-level mitochondrial consensus sequences using annotated coordinates and translated using the vertebrate mitochondrial genetic code. Protein sequences were aligned using MAFFT, and the resulting alignments were back-translated to codon alignments using PAL2NAL v14.1 (Suyama et al. 2006).

### Estimation of d_N_, d_S_, and d_N_/d_S_

Gene-level nonsynonymous divergence (d_N_), synonymous divergence (d_S_), and the ratio d_N_/d_S_ were estimated using codeml in PAML v4.10.10 under the one-ratio M0 model (Yang 2007). All genes were analyzed using a common unrooted topology of the 27 study populations, with branch lengths estimated independently for each gene. Analyses used seqtype=1, runmode=0, model=0, NSsites=0, CodonFreq=2, and cleandata=1, with the standard genetic code for nuclear genes and the vertebrate mitochondrial genetic code for mitochondrial genes.

Codeml likelihoods, transition/transversion ratios, tree lengths, and rate estimates were retained for quality control. Failed runs, invalid estimates, and nonfinite focal statistics were excluded. Invariant nuclear genes were retained with d_N_ = d_S_ = 0, whereas their d_N_/d_S_ values were treated as undefined. Primary nuclear d_N_/d_S_ comparisons required d_S_ > 0, at least one expected synonymous substitution, and a finite ratio below the upper optimization boundary. A stricter threshold of at least two expected synonymous substitutions was examined as a sensitivity analysis. Genes with identifiable synonymous divergence but no nonsynonymous divergence were assigned d_N_/d_S_ = 0. Mitochondrial d_N_/d_S_ comparisons required d_S_ ≥0.001 and an estimated ratio <999. Genes excluded from ratio comparisons remained eligible for d_N_ or d_S_ comparisons when those statistics were identifiable. No pseudocount was added to d_S_, and d_N_, d_S_, and d_N_/d_S_ were interpreted jointly.

### Estimation of π_N_, π_S_, and π_N_/π_S_

Within-population π_N_ and π_S_ were calculated from synchronized Pool-seq allele counts. Each coding position was assigned to its representative transcript using exon order, strand, and codon phase. All possible single-nucleotide changes at callable codons were classified as synonymous or nonsynonymous under the appropriate nuclear or mitochondrial genetic code, providing gene-specific synonymous and nonsynonymous site counts.

At each position, nucleotide diversity was calculated as:

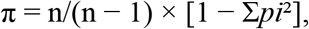

where *pi* is the frequency of allele i and n is the number of sampled chromosomes. Nuclear pools used 2N chromosomes and mitochondrial pools used N mitochondrial genomes. Diversity attributable to nonsynonymous and synonymous changes was summed and divided by the corresponding numbers of mutational opportunities to obtain π_N_ and π_S_. Ambiguous codons, start and stop codons, positions near indels, and insufficiently covered sites were excluded.

Gene-level π_N_ and π_S_ were averaged across eligible populations, and π_N_/π_S_ was calculated as the ratio of the across-population means. This avoided disproportionate effects from individual populations with very small π_S_. Functional comparisons and multiple-testing correction followed the divergence analyses.

### Functional comparisons and statistical analyses

Divergence and polymorphism statistics were compared across six predefined, complementary functional classifications (Tables S3 and S4; Fig. 2 and Figs. S2–S5). These classifications addressed different biological questions and were not mutually exclusive across panels.

We compared mitochondrial OXPHOS genes, nuclear OXPHOS structural subunits, and nuclear OXPHOS assembly factors to assess differences among mitochondrially encoded respiratory-chain components, nuclear-encoded structural components, and proteins involved primarily in complex assembly. These comparisons were also conducted separately by OXPHOS complex. Nmt-RPs and Nmt-ARSs were compared with their respective cytosolic counterparts to assess differences between mitochondrial and cytosolic translation systems. Mitochondrial OXPHOS genes were additionally compared with nuclear core and noncore OXPHOS subunits. Finally, direct n-mt, indirect n-mt, and non-n-mt genes were compared at the genome-wide level. Predefined pairwise comparisons used two-sided Wilcoxon rank-sum tests. Where multiple pairwise comparisons were performed, *P* values were adjusted using the Benjamini–Hochberg procedure separately within each functional classification and evolutionary statistic. Genome-wide medians calculated from quality-filtered nuclear genes were included as descriptive reference values rather than as independent statistical groups. Genes with undefined d_N_/d_S_ or π_N_/π_S_ were excluded from the corresponding ratio comparisons but retained in comparisons of individual component statistics when those estimates remained identifiable.

### Evolutionary rate covariation

Evolutionary rate covariation (ERC) was used to test whether mitochondrial and nuclear evolutionary rates changed coordinately across corresponding branches of the population phylogeny. In contrast to d_N_/d_S_, which measures nonsynonymous divergence relative to synonymous divergence, ERC evaluates whether the same lineages exhibit relative acceleration or deceleration in different genes. Analyses were conducted at the individual-gene, functional gene-set, and OXPHOS-complex levels.

ERC calculations used codeml total branch lengths estimated independently for 15,241 nuclear genes and all 13 mitochondrial protein-coding genes. All genes were analyzed on the same fixed, unrooted 27-population topology, producing 51 corresponding branches. The topology was fixed to permit direct matching of branch identities, whereas branch lengths were estimated separately from each alignment.

Raw branch lengths reflect branch duration and shared evolutionary history in addition to gene-specific rate variation. For each branch *b*, an empirical background rate, *B*_b_, was therefore calculated as the mean branch length across the 15,241 retained nuclear genes. The mean was used because the large number of genes with zero estimated change on individual branches caused many branch-specific medians to equal zero.

For each gene *g*, observed branch lengths were regressed against the genome-wide background using a through-origin model:

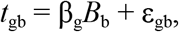

where *t*_gb_ is the estimated length of branch *b* for gene *g*, β_g_ represents the overall evolutionary rate of the gene relative to the nuclear background, and ε_gb_ represents its branch-specific deviation from expectation. β_g_ was constrained to be nonnegative. Residuals were centered and divided by their root-mean-square value to obtain standardized relative evolutionary rates. The same nuclear background and normalization procedure were applied to both nuclear and mitochondrial genes.

The primary mitochondrial predictor was the branch-specific median standardized rate across all 13 mitochondrial protein-coding genes. Spearman correlations were calculated between this mitochondrial composite and the 51-branch relative-rate vector of each nuclear gene. For each gene, we retained the correlation coefficient, nominal *P* value, number of matched branches, and Benjamini–Hochberg-adjusted *P* value. Additional analyses used individual mitochondrial genes and mitochondrial composites corresponding to Complexes I, III, IV, and V.

The distribution of gene-level ERC coefficients among non-n-mt genes provided the empirical reference for threshold and enrichment analyses. Genes exceeding the 99th percentile of this reference distribution were classified as high-covariation candidates; the 95th and 97.5th percentiles were examined as sensitivity thresholds. Enrichment of mitochondrial functional categories above these thresholds was evaluated relative to non-n-mt genes using one-sided Fisher’s exact tests. Full ERC distributions were compared with the non-n-mt distribution using one-sided Wilcoxon rank-sum tests for elevated correlations, with Cliff’s delta reported as an effect-size measure. Benjamini–Hochberg correction was applied across category-level comparisons. All 15,241 nuclear genes retained in the ERC analysis were additionally summarized as a descriptive genome-wide reference, but were not treated as an independent comparator for enrichment tests.

Functional gene-set ERC followed the bootstrap framework of Weaver et al. (2022). For each branch, nuclear and mitochondrial composite rates were calculated as the arithmetic means of standardized rates across genes in their respective sets. Spearman correlation was calculated between the resulting nuclear and mitochondrial composite-rate vectors. This gene-set analysis therefore differed from the individual-gene screening, which used the median mitochondrial standardized rate as its predictor. Uncertainty was evaluated using 10,000 bootstrap replicates in which nuclear and mitochondrial genes were independently sampled with replacement from their respective sets, and both composite-rate vectors were recalculated in each replicate. A positive gene-set association was considered supported when its percentile-based 95% bootstrap confidence interval excluded zero. Differences between mitochondrial and cytosolic translation-gene sets were summarized using distributions of bootstrap ERC differences.

Within-complex analyses compared nuclear and mitochondrial genes belonging to the same respiratory complex. Nuclear Complex I genes were compared with ND1–ND6 and ND4L; nuclear Complex III genes with CYTB; nuclear Complex IV genes with COX1–3; and nuclear Complex V genes with ATP6 and ATP8. Because Complex II is entirely nuclear encoded, its subunits were compared with the composite of all 13 mitochondrial protein-coding genes as a pathway-level comparison rather than a direct within-complex mitonuclear test. Each association was evaluated using 10,000 gene-bootstrap replicates.

The strongest within-complex association was subjected to leave-one-out analyses by independently removing each mitochondrial gene, nuclear gene, and phylogenetic branch. Branch-label permutations further tested whether the association depended on matching nuclear and mitochondrial rates from corresponding branches. Mitochondrial rates were either permuted among all 51 branches or separately within terminal and internal branches. Each test used 100,000 permutations. Upper-tail empirical probabilities were calculated as

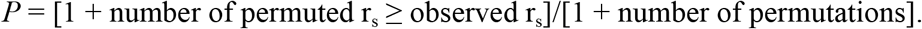

Both unrestricted and terminal/internal-branch-stratified permutation tests were applied to all five OXPHOS complexes, with Benjamini–Hochberg correction across complexes performed separately for each permutation scheme.

### Population branch statistic analyses

The population branch statistic (PBS) was used to quantify freshwater-specific mitochondrial and nuclear differentiation, with a Norwegian stickleback population serving as the outgroup (Yi et al. 2010). Pairwise *F*_ST_ was estimated from allele counts using grenedalf and the filters described above. Mitochondrial estimates excluded the control region; nuclear estimates were calculated for annotated gene intervals and chromosome XXI.

For each freshwater population A, its corresponding regional anadromous reference B, and the Norway outgroup C, pairwise *F*_ST_ values were transformed as

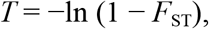

and freshwater-branch PBS was calculated as

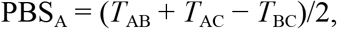

RS and SAY served as the regional anadromous references for Alaska and British Columbia, respectively. The population comparisons and calculation of freshwater-branch PBS are illustrated in Fig. S6.

Unadjusted mitonuclear concordance was summarized using Pearson correlations between mitochondrial PBS and gene-level nuclear PBS across freshwater populations. Nuclear OXPHOS correlations used the 19 long-term freshwater populations remaining after exclusion of AMO. Other mitochondrial functional classes and cytosolic comparison groups were also examined to assess whether positive associations were specific to genes with mitochondrial functions.

Among the 70 nuclear OXPHOS genes assigned to complexes I–V, differences in correlation strength were evaluated using the Kruskal–Wallis statistic, with significance assessed by 9,999 permutations of gene-to-complex assignments while preserving complex sizes. Differences among complexes in the proportion of positively correlated genes were evaluated using Fisher’s exact test with 9,999 Monte Carlo replicates. Positive correlations were defined as r > 0 regardless of individual-gene significance. Each complex was additionally tested for enrichment of positive correlations relative to the remaining complexes using a one-sided Fisher’s exact test, with Benjamini–Hochberg correction across the five comparisons.

For each of the 72 nuclear OXPHOS genes, we subsequently tested whether its association with mitochondrial PBS persisted after accounting for region and background nuclear differentiation: nuclear PBS ∼ mitochondrial PBS + chromosome XXI PBS + region.

These models used the same 19 long-term freshwater populations as the unadjusted OXPHOS correlations. We recorded the mitochondrial PBS coefficient and partial R² for each gene. For visualization, residual nuclear and mitochondrial PBS values were obtained by independently regressing each measure on chromosome XXI PBS and region.

Significance was evaluated using 10,000 Freedman–Lane permutations. Residuals from the reduced model containing chromosome XXI PBS and region were permuted within the region, added to the fitted reduced-model values, and used to refit the full model. Two-sided empirical *P* values were calculated from the absolute mitochondrial PBS coefficients using a plus-one correction. Within the predefined nuclear OXPHOS set, genes with positive mitochondrial PBS coefficients and nominal permutation *P* < 0.05 were considered candidate positive associations. Benjamini–Hochberg-adjusted *P* values across the 72 genes were additionally reported in Table S6.

### Regional allele-frequency change and parallelism

Nuclear allele frequencies were calculated from A, T, C, and G read counts, requiring depth ≥20. At each SNP, the nucleotide with the largest total count across retained populations was defined as the focal allele and used consistently in both regions. Signed allele-frequency change was calculated as

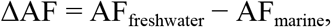

using RS and SAY as the marine references for Alaska and British Columbia, respectively.

Regional comparisons were restricted to the 72 curated nuclear OXPHOS genes and included all 25 freshwater populations: 11 from Alaska and 14 from British Columbia. One representative SNP was selected per gene so that both regions were compared at the same genomic position and for the same focal allele. Eligible SNPs required complete coverage across all freshwater populations and their corresponding regional marine references.

Regional median ΔAF values were calculated across all freshwater populations within each region. For genes containing SNPs with concordant, nonzero regional median ΔAF values, selection was restricted to those SNPs. Eligible SNPs were ranked by the smaller of the two absolute regional median ΔAF values, prioritizing variants with strong effects in both regions. Ties were resolved sequentially by the mean absolute regional median ΔAF, the minimum within-region directional concordance, the minimum regional mean sequencing depth, and genomic position. If no SNP showed concordant regional directions, the same ranking criteria were applied to all eligible shared SNPs.

Regional correspondence was summarized using the Pearson correlation between Alaska and British Columbia median ΔAF values and the number of genes with concordant directions. Because representative-SNP selection favored directional concordance, these summaries were interpreted descriptively rather than as independent tests of parallelism.

Genes with |median ΔAF| ≥0.5 in at least one region were designated high-effect candidates, yielding ten genes for the population-level frequency plots. These plots distinguished long-term freshwater populations from the five recently colonized populations. Black segments indicated median frequencies among long-term freshwater populations, whereas regional ΔAF estimates and candidate thresholds were calculated using all freshwater populations, including recently colonized populations.

### Linkage disequilibrium and SNP-level association models

Read-backed LD was estimated separately by population and OXPHOS gene using LDx (Feder et al. 2012). SNPs required mapping and base qualities ≥30 and depth ≥20. SNP pairs with *R*² ≥0.2 were connected into LD groups. Within each group, the SNP with the largest absolute ΔAF was retained for downstream association tests.

Geographic and mitochondrial-lineage association analyses included the same 19 long-term freshwater populations, excluding AMO and the five recently colonized populations. Geographic associations were evaluated using quasibinomial generalized linear models with a logit link, fitted to focal- and alternative-allele read counts:

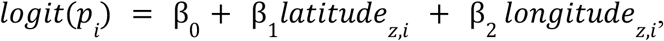

where *p_i_* is the expected focal-allele frequency in population *i*, and *latitude_z_*_,*i*_ and *longitude_z_*_,*i*_ are standardized latitude and longitude, respectively. The full model was compared with an intercept-only model using an F test. Effect size was defined as the range of fitted allele frequencies across the observed geographic coordinates.

Mitochondrial-lineage associations were evaluated using quasibinomial generalized linear models fitted to the same paired allele counts:

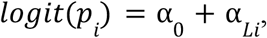

where *L_i_* denotes the mitochondrial lineage of population *i*, and α*L*_i_ represents the corresponding lineage effect, with the reference-lineage effect set to zero. Each model was compared with an intercept-only model using an F test. Effect size was defined as the maximum difference in depth-weighted allele frequencies among mitochondrial lineages. Benjamini–Hochberg correction was applied separately across estimable SNP tests in each association analysis. Candidate SNPs were required to satisfy both *q* < 0.05 and an effect size ≥0.20.

Gene-level lineage dependence was evaluated using population profiles of signed ΔAF across all LD-pruned SNPs within each OXPHOS gene. Pairwise dissimilarity was calculated as

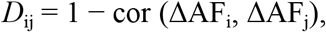

where ΔAF_i_ and ΔAF_j_ are vectors of signed allele-frequency changes at shared SNPs in populations i and j. The contrast between within- and between-lineage distances was summarized as

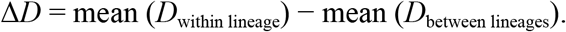

Negative ΔD indicates greater similarity of nuclear allele-frequency profiles among populations sharing a mitochondrial lineage. One-sided significance was evaluated by permuting lineage labels while retaining the observed distance matrix. Analyses were conducted across all 19 populations and separately within Alaska and British Columbia. Nominal permutation *P* < 0.05 was treated as suggestive evidence, and Benjamini–Hochberg-adjusted *q*-values were reported across genes separately for the combined and region-specific analyses (Table S9).

### Identification of recurrent nonsynonymous variants

Nonsynonymous variation was evaluated across nuclear loci representing the 72 curated OXPHOS subunit symbols. Analyses included the 20 long-term freshwater populations, including AMO, and the regional anadromous references RS and SAY. Representative CDS models were reconstructed in transcript orientation using strand-aware exon order and CDS phase. Coding SNPs were translated using the standard nuclear genetic code. Synonymous variants and codons containing ambiguous bases, gaps, incomplete coverage, or stop codons were excluded. Because Pool-seq data do not resolve codon-level phase, multiple SNPs within the same codon were evaluated individually rather than combined into inferred codon haplotypes.

Alternative-amino-acid frequencies were calculated from nucleotide counts when both a freshwater population and its regional anadromous reference had depth ≥20. Population-specific frequency differences were calculated as

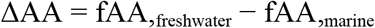

Regional summaries required valid ΔAA estimates for at least three freshwater populations. Regional effect sizes were summarized using median ΔAA. Marine-associated and freshwater-associated amino-acid states were identified from the observed population frequencies, independently of the reference and alternative labels in the genome assembly; these labels were not interpreted as ancestral and derived states.

Variants were matched between regions by gene, amino-acid position, reference residue, and alternative residue. Exact two-sided sign-flip tests were performed within each region using the absolute mean population-specific ΔAA as the test statistic. Suggestive parallel variants required concordant regional directions, |median ΔAA| ≥0.20, and nominal *P* < 0.05 in both regions. Stringent parallel candidates additionally required |median ΔAA| ≥0.50 in both regions. These nominal tests were used to prioritize candidates rather than establish genome-wide significance. Biochemical similarity was summarized using BLOSUM62 scores (Henikoff and Henikoff 1992). Functional tolerance was predicted using SIFT4G v2.0.0 (Ng and Henikoff 2003; Vaser et al. 2016) with the corresponding regional marine-reference protein sequence as the query background, thereby evaluating amino-acid changes relative to the marine-associated sequence. Scores ≤0.05 were classified as deleterious and scores >0.05 as tolerated. These predictions were used for candidate prioritization rather than as independent evidence of adaptation.

### COX4I1 sequence conservation and Complex IV structural predictions

To examine whether the three recurrent freshwater-associated COX4I1 substitutions could alter a mitonuclear interface, we reconstructed two full-length 169-amino-acid COX4I1 sequences: the predominant marine-associated A58–F124–F133 state and the freshwater-associated V58–Y124–L133 state. These sequences represent combinations of population-major amino-acid states inferred from Pool-seq frequencies and should not be interpreted as experimentally phased individual haplotypes.

For cross-species comparisons, the two reconstructed stickleback sequences were aligned with COX4I1 protein sequences from eight other vertebrate species: zebrafish (*Danio rerio*, NP_999866.1), medaka (*Oryzias latipes*, XP_004067266.1), blue catfish (*Ictalurus furcatus*, XP_053478496.1), spotted gar (*Lepisosteus oculatus*, XP_015223741.1), western clawed frog (*Xenopus tropicalis*, XP_012816550.1), chicken (*Gallus gallus*, NP_001025748.1), human (*Homo sapiens*, NP_001852.1), and mouse (*Mus musculus*, UniProt P19783, sequence version 2). Sequences were retrieved from NCBI Protein and aligned using Clustal Omega v1.2.4 with default settings (Sievers et al. 2011). Alignment columns corresponding to stickleback residues 58, 124, and 133 were identified using the ungapped stickleback sequence as the numbering reference. Conservation was assessed descriptively by comparing amino-acid identities and physicochemical properties at these positions. Local alignment windows encompassing each focal site and five alignment columns on either side were visualized with residues colored by physicochemical class.

For structural predictions, COX4I1 was modeled together with the three mitochondrially encoded catalytic subunits of Complex IV: MT-CO1, MT-CO2, and MT-CO3. Mitochondrial protein sequences were translated from population-consensus coding sequences. MT-CO2 and MT-CO3 were invariant at the amino-acid level among sampled populations. The only MT-CO1 amino-acid polymorphism, I83V, was restricted to AMO. The common MT-CO1 sequence was consequently used in both modeled complexes to isolate structural differences associated with the COX4I1 sequence states.

Protein complexes were predicted using ColabFold v1.6.1 with AlphaFold-Multimer v3, MMseqs2-generated multiple-sequence alignments, three recycles, no templates, and no Amber relaxation (Evans et al. 2021; Jumper et al. 2021; Mirdita et al. 2022). Five models were generated for each COX4I1 sequence state and ranked according to the ColabFold multimer confidence score. Corresponding marine- and freshwater-associated models were compared. Prediction confidence was assessed using predicted local distance difference test scores (pLDDT), predicted TM scores (pTM), interface predicted TM scores (ipTM), and predicted aligned error (PAE).

Marine- and freshwater-associated models were superimposed using the mitochondrially encoded Complex IV core. For COX4I1 residues 58, 124, and 133, the minimum distance between any residue heavy atom and any heavy atom in each mitochondrial subunit was calculated across the five models for each sequence state. A minimum distance ≤5 Å was used as an operational criterion for a predicted intersubunit contact (Weaver et al. 2022); distances >5 Å and ≤10 Å were classified as interface-proximal. Structural predictions were used to compare overall complex architecture and local intersubunit geometry, rather than to establish biochemical or fitness effects. Models were superimposed, inspected, and visualized using UCSF ChimeraX (Pettersen et al. 2021).

## Data and resource availability

The sequencing data analyzed in this study were obtained from Weber et al. (2022) and are available through the repositories listed in that publication. The analysis code and scripts used to generate the results and figures are available on GitHub at: https://github.com/yuchangyue1209/Mt-nuclear-co-evolution-paper/tree/main/publication Additional processed data and supporting files are available from the corresponding author upon request.

### Acknowledgements

This work was supported by the National Institutes of Health (NIAID 1R01AI123659 to D.I.B.; NIGMS 1R35GM142891 to J.N.W.) and by startup funding from the University of Wisconsin-Madison (J.N.W.). We thank Dr. Daniel Jeffries for insightful discussion on mitochondrial genetic variation in stickleback. We also thank Dr. John Pool for helpful discussion of the PBS analysis. During the preparation of this work, the authors used ChatGPT (OpenAI) and Claude (Anthropic) to assist with language editing and to organize and debug the analysis code. The primary research questions were created solely by the authors. The authors reviewed and verified all AI-assisted outputs and take full responsibility for the manuscript.

## Author Contributions

C.Y. and J.N.W. conceived the study. C.Y. performed the analyses. C.Y. and J.N.W. interpreted the results and wrote the manuscript. C.Y. visualized the results. D.I.B. acquired funding to generate the raw data and advised on methodology and writing. J.N.W. acquired funding and supervised the work. All authors reviewed and approved the final manuscript.

## Conflict of Interest Disclosure

The authors declare no conflict of interest.

## Notes

https://github.com/yuchangyue1209/Mt-nuclear-co-evolution-paper/tree/main/publication

### Competing Interest Statement

The authors have declared no competing interest.

### Summary of Updates

We revised the equation formatting in the Methods section and updated the fonts in Figure 1.

