## Supplementary Information (Figures S1 to S9) for "Parallel and lineage-dependent mitonuclear evolution in threespine stickleback"

### Contents

#### Supplementary Figures

**Figure S1.** Sampling locations of threespine stickleback populations in Alaska and British Columbia.

**Figure S2.** Patterns of within-population nonsynonymous and synonymous nucleotide diversity across OXPHOS, ribosomal protein, and aminoacyl-tRNA synthetase genes.

**Figure S3.** Divergence and within-population nucleotide diversity across OXPHOS complexes and gene classes.

**Figure S4.** Divergence and within-population nucleotide diversity across mitochondrial OXPHOS and nuclear core and noncore OXPHOS subunit genes.

**Figure S5.** Divergence and within-population nucleotide diversity across direct n-mt, indirect n-mt, and non-n-mt genes.

**Figure S6.** Calculation of population branch statistics for freshwater stickleback populations.

**Figure S7.** Concordance between nuclear and mitochondrial PBS across additional mitochondrial-associated and cytosolic gene classes.

**Figure S8.** Linkage disequilibrium structure across nuclear OXPHOS genes and freshwater populations.

**Figure S9.** Region-specific mitochondrial-lineage similarity in nuclear OXPHOS allele-frequency profiles.

### Supplementary Tables

Complete tables are provided as separate Excel files (TableS1.xlsx–TableS10.xlsx).

**Table S1.** Population metadata and analysis inclusion.

**Table S2.** Curated gene sets, orthology assignments, and analysis inclusion.

**Table S3.** Summary of coding-sequence divergence across mitochondrial-associated gene classes.

**Table S4.** Summary of nucleotide diversity across mitochondrial-associated gene classes.

**Table S5.** Summary of evolutionary-rate covariation across gene sets and OXPHOS complexes.

**Table S6.** Nuclear OXPHOS differentiation and unadjusted and demographic-adjusted mitonuclear PBS associations.

**Table S7.** Regional freshwater–marine allele-frequency shifts at representative SNPs in nuclear OXPHOS genes.

**Table S8.** Geographic and mitochondrial-lineage associations of nuclear OXPHOS allele frequencies.

**Table S9.** Within- and between-mitochondrial-lineage similarity of nuclear OXPHOS allele-frequency profiles.

**Table S10.** Nonsynonymous variants in nuclear OXPHOS genes and regional freshwater–marine comparisons.

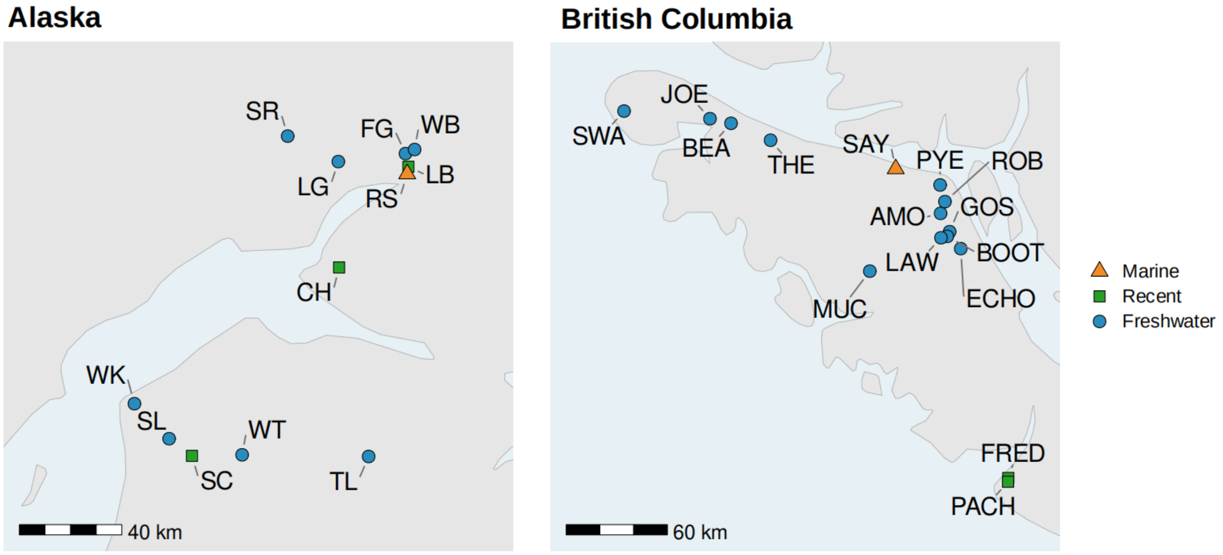


**Figure S1. Sampling locations of threespine stickleback populations in Alaska and British Columbia.** The dataset comprised 20 established freshwater populations (blue circles), five recently colonized freshwater populations (green squares), and two regional marine reference populations (orange triangles). Light blue indicates water and gray indicates land. Population abbreviations and sampling information are provided in Table S1.

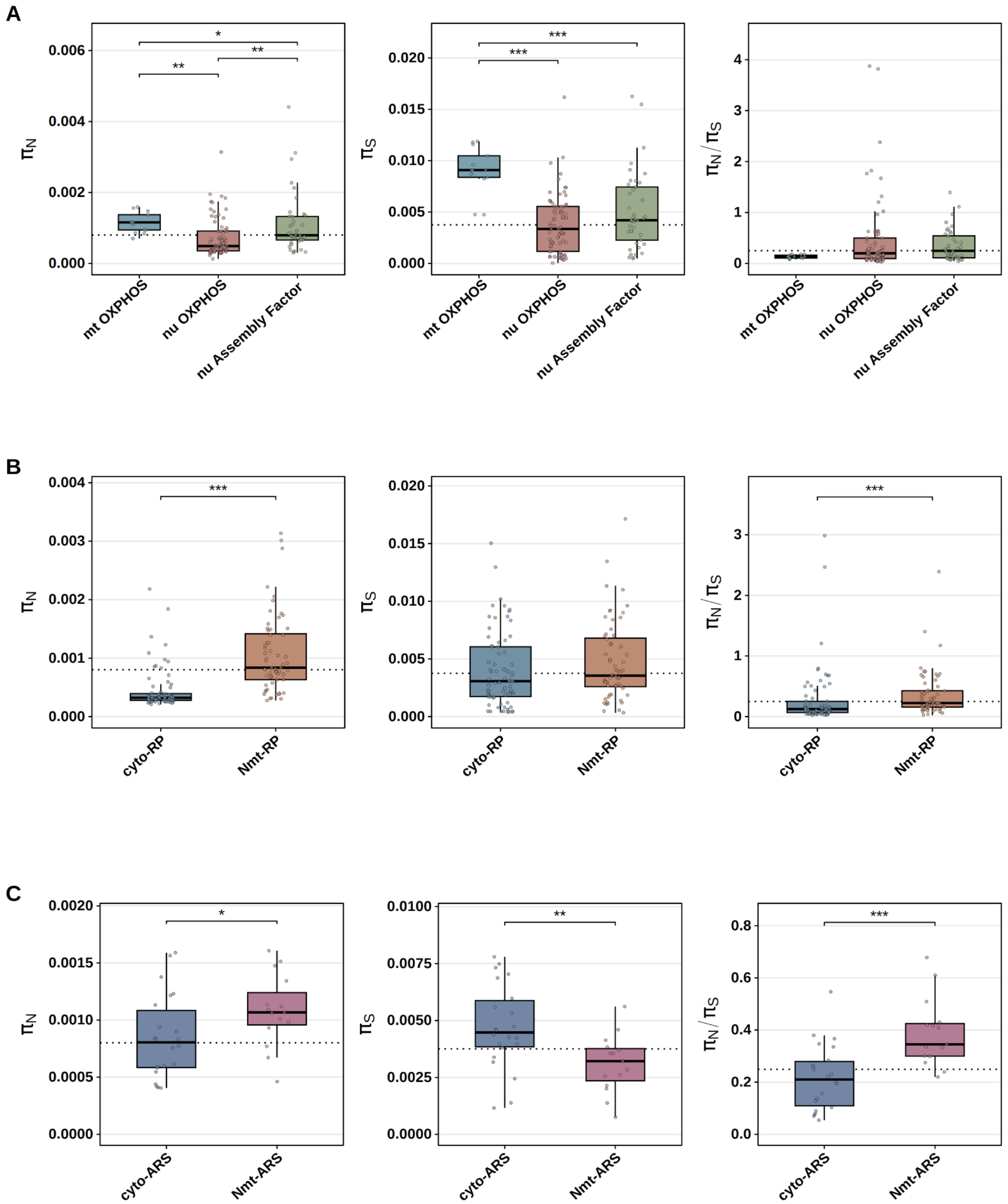

A
B
C
πN
πS
πN/πS
mt OXPHOS
nu OXPHOS
nu Assembly Factor
cyto-RP
Nmt-RP
cyto-ARS
Nmt-ARS

**Figure S2. Patterns of within-population nonsynonymous and synonymous nucleotide diversity across OXPHOS, ribosomal protein, and aminoacyl-tRNA synthetase genes.** Distributions of $\pi_N$, $\pi_S$, and $\pi_N/\pi_S$ are shown for (A) mitochondrial OXPHOS genes (mt OXPHOS), nuclear OXPHOS subunit genes (nu OXPHOS), and nuclear OXPHOS assembly-factor genes; (B) cytosolic and nuclear-encoded mitochondrial ribosomal proteins (RPs); and (C) cytosolic and nuclear-encoded mitochondrial aminoacyl-tRNA synthetases (ARSs); boxes indicate medians and interquartile ranges, whiskers extend to the most extreme values within 1.5 times the interquartile range, and points represent individual genes. Dotted horizontal lines indicate genome-wide medians across all quality-eligible nuclear genes. Pairwise differences were evaluated using two-sided Wilcoxon rank-sum tests with Benjamini–Hochberg correction where multiple comparisons were performed. Asterisks denote adjusted significance levels: $P < 0.05$ (*), $P < 0.01$ (**), $P < 0.001$ (***).

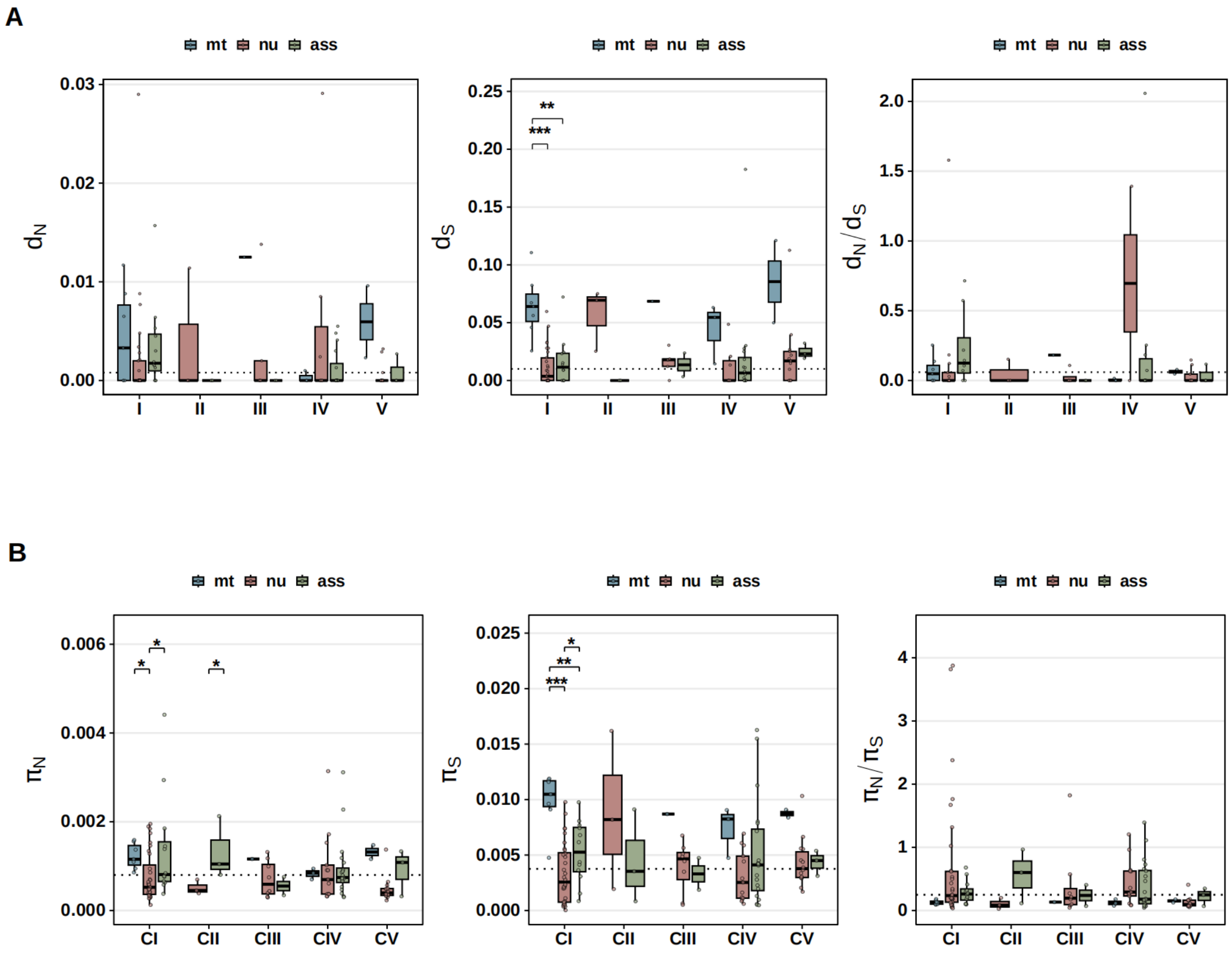


**Figure S3. Divergence and within-population nucleotide diversity across OXPHOS complexes and gene classes.** Distributions of (A) $d_N$, $d_S$, and $d_N/d_S$ and (B) $\pi_N$, $\pi_S$, and $\pi_N/\pi_S$ are shown for OXPHOS complexes I–V. Colors distinguish mitochondrial OXPHOS subunit genes (mt), nuclear OXPHOS subunit genes (nu), and nuclear OXPHOS assembly-factor genes (ass). Boxes indicate medians and interquartile ranges; whiskers extend to the most extreme values within 1.5 times the interquartile range. Each point represents an individual gene. Dotted horizontal lines indicate genome-wide nuclear medians calculated from genes with eligible estimates for each metric. Pairwise differences were evaluated using two-sided Wilcoxon rank-sum tests with Benjamini–Hochberg correction. Asterisks denote adjusted significance levels $P < 0.05$ (*), $P < 0.01$ (**), $P < 0.001$ (***).

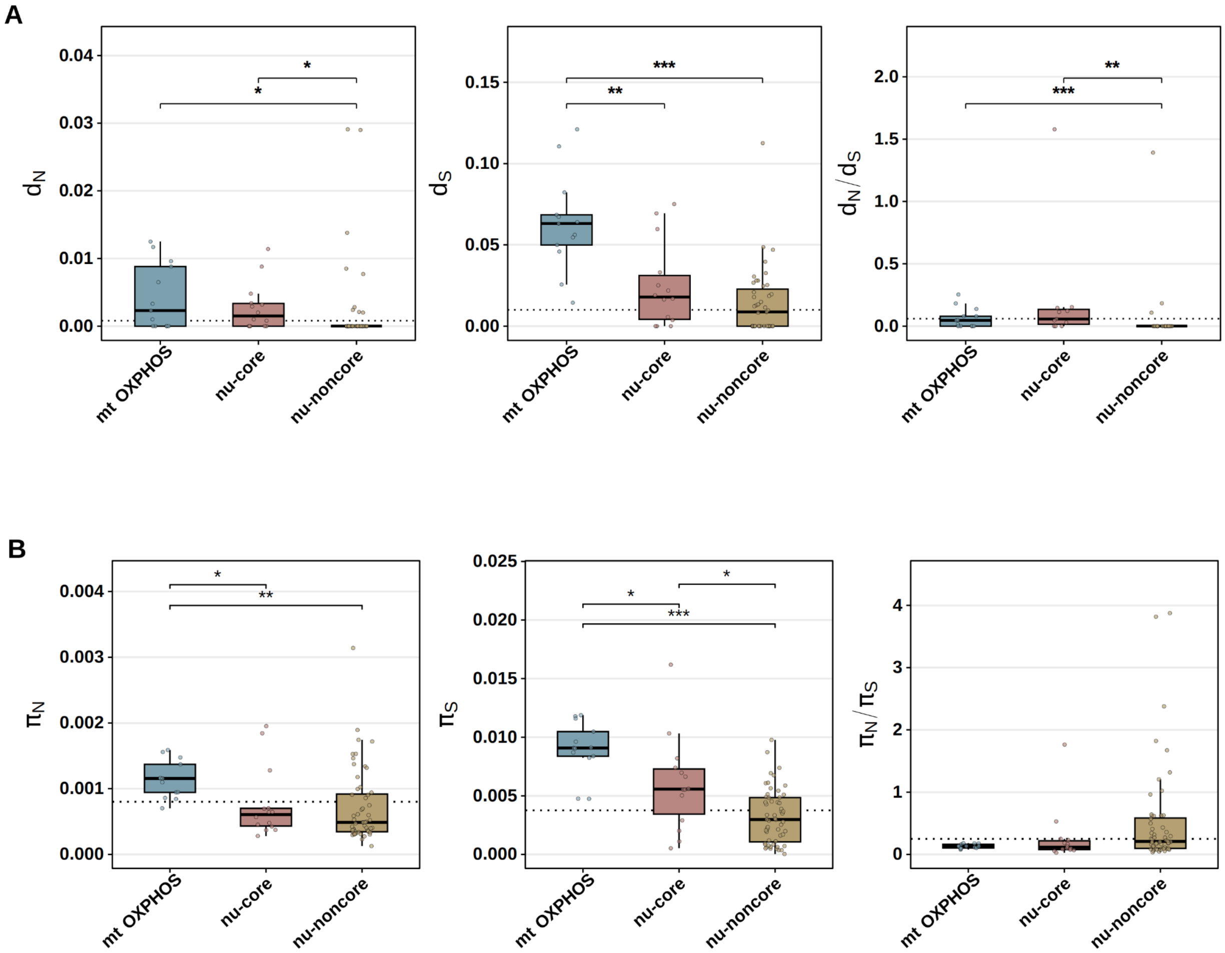


**Figure S4. Divergence and within-population nucleotide diversity across mitochondrial OXPHOS and nuclear core and noncore OXPHOS subunit genes.** Distributions of (A) $d_N$, $d_S$, and $d_N/d_S$ and (B) $\pi_N$, $\pi_S$, and $\pi_N/\pi_S$ are shown for mitochondrial OXPHOS genes (mt OXPHOS), nuclear core OXPHOS subunit genes (nu-core), and nuclear noncore OXPHOS subunit genes (nu-noncore). Boxes indicate medians and interquartile ranges; whiskers extend to the most extreme values within 1.5 times the interquartile range. Each point represents an individual gene. Dotted horizontal lines indicate genome-wide nuclear medians calculated from genes with eligible estimates for each metric. Pairwise differences were evaluated using two-sided Wilcoxon rank-sum tests with Benjamini–Hochberg correction. Asterisks denote adjusted significance levels $P < 0.05$ (*), $P < 0.01$ (**), $P < 0.001$ (***).

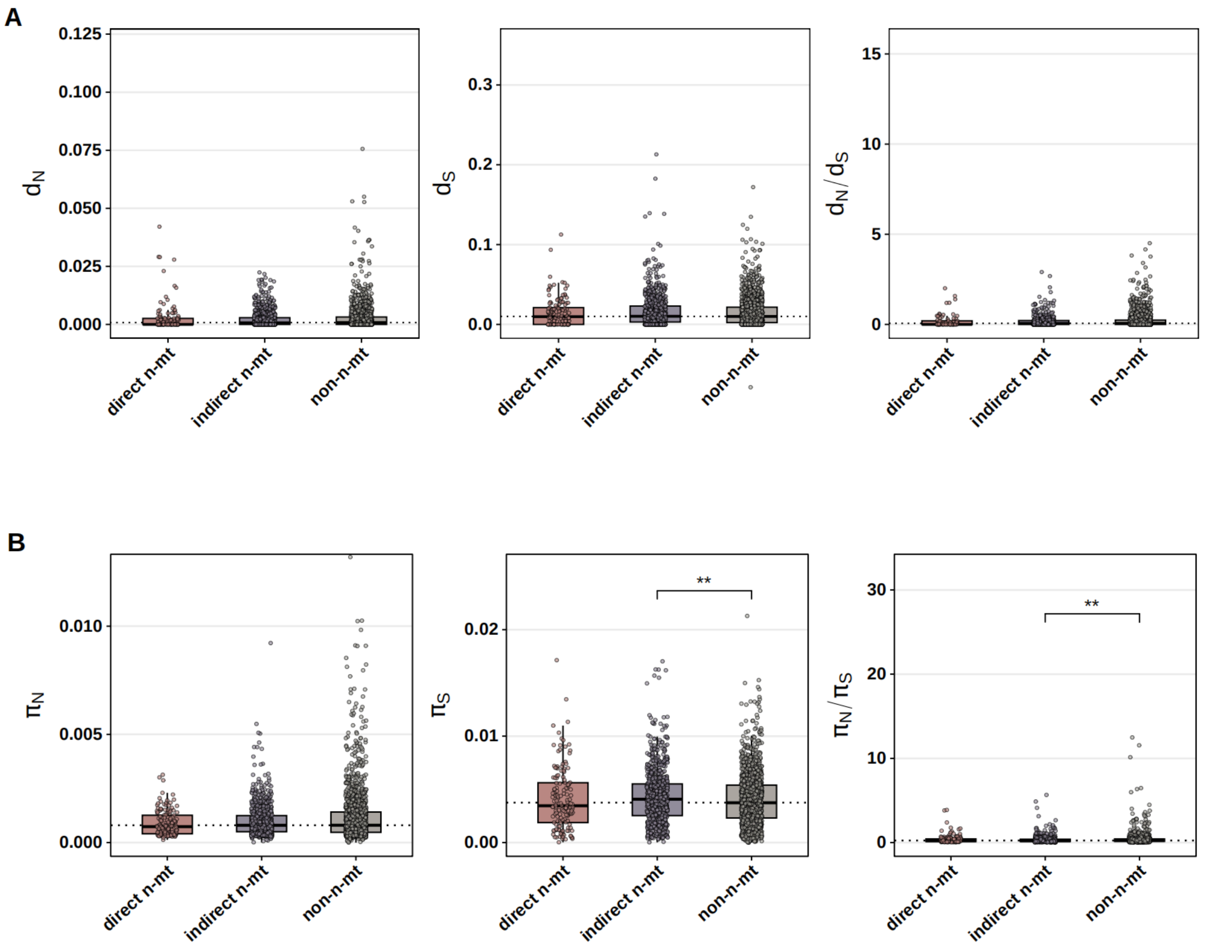


**Figure S5. Divergence and within-population nucleotide diversity across direct n-mt, indirect n-mt, and non-n-mt genes.** Distributions of (A) $d_N$, $d_S$, and $d_N/d_S$ and (B) $\pi_N$, $\pi_S$, and $\pi_N/\pi_S$ are shown for the three nuclear gene classes. Boxes indicate medians and interquartile ranges; whiskers extend to the most extreme values within 1.5 times the interquartile range. Points represent individual genes and were subsampled for visualization; boxplots and statistical tests used all eligible genes. Dotted horizontal lines indicate genome-wide nuclear medians calculated from genes with eligible estimates for each metric. Pairwise differences were evaluated using two-sided Wilcoxon rank-sum tests with Benjamini–Hochberg correction. Asterisks denote adjusted significance levels $P < 0.05$ (*), $P < 0.01$ (**), $P < 0.001$ (***).

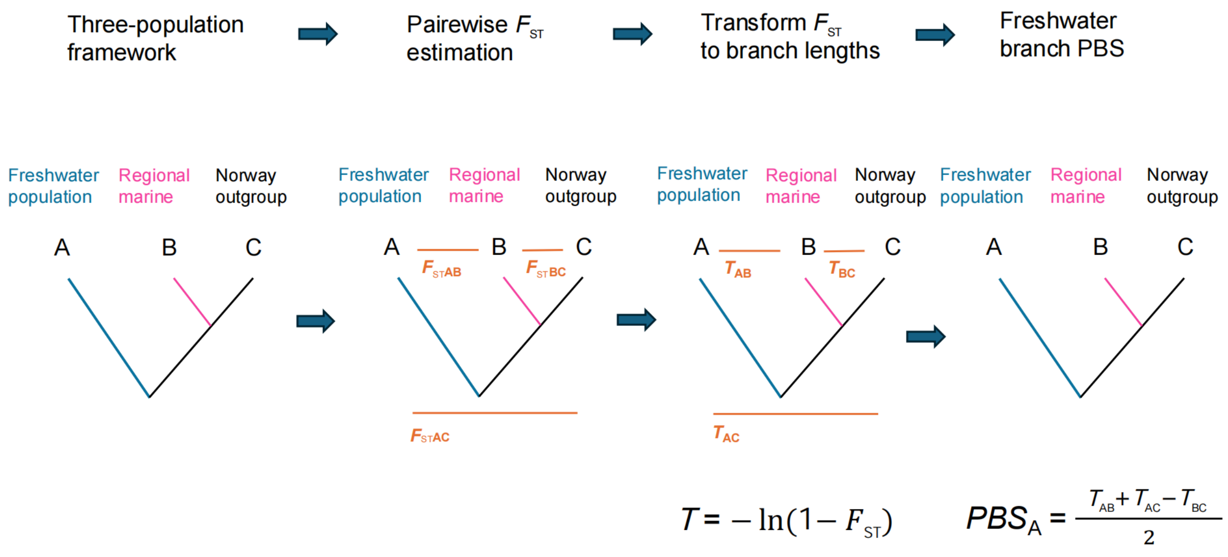


**Figure S6. Calculation of population branch statistics for freshwater stickleback populations.** For each focal freshwater population (A), PBS was calculated relative to the corresponding regional marine reference population (B) and the Norway outgroup population (C). Pairwise *F*st values were estimated for the three population pairs and transformed to branch-length distances as $T = -\ln(1 - F_{ST})$. The freshwater-specific branch statistic was then calculated as $\text{PBS}_A = (T_{AB} + T_{AC} - T_{BC})/2$, which represents differentiation specific to the focal freshwater population after accounting for divergence between the marine reference and the outgroup.

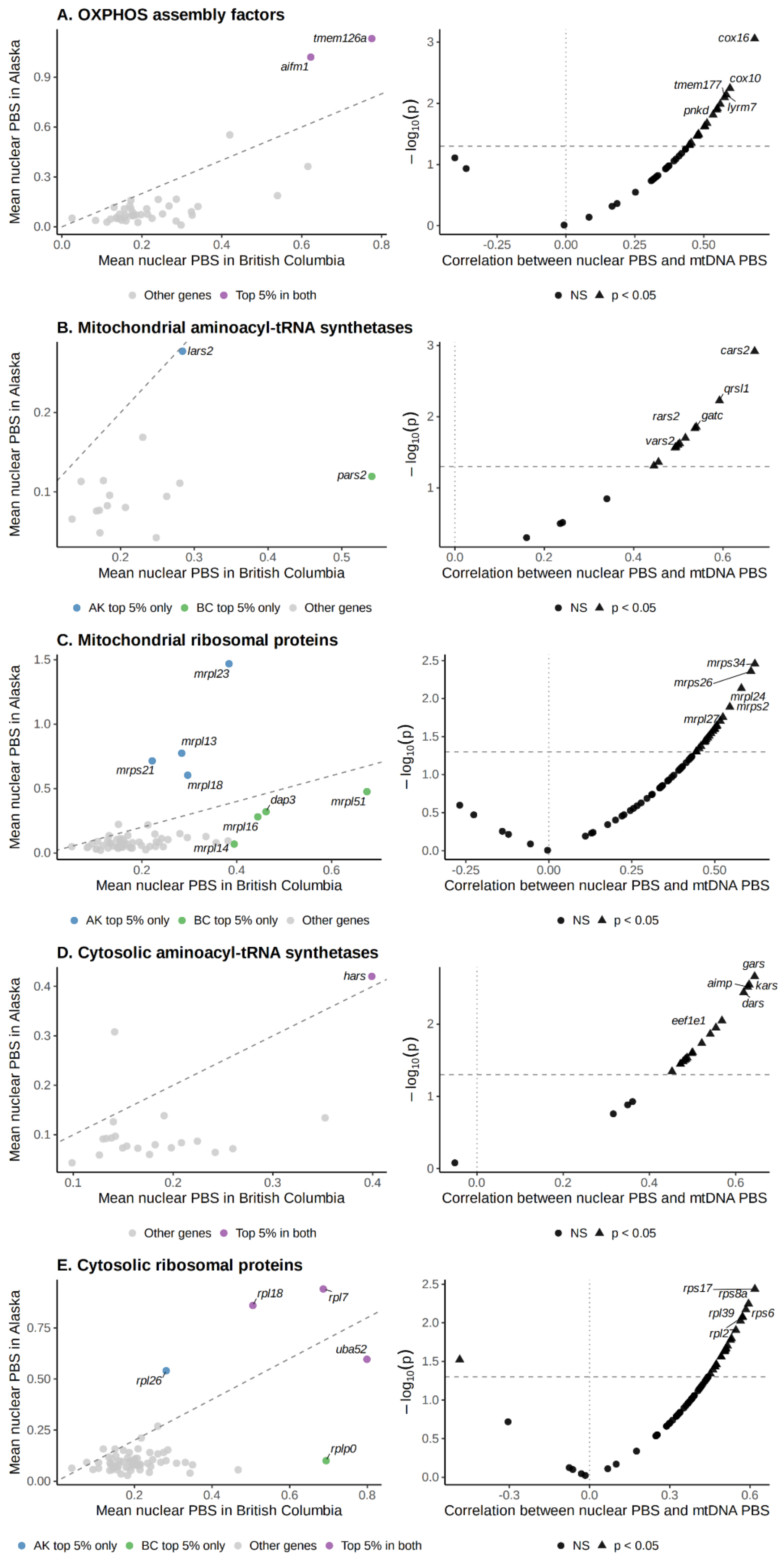
A. OXPHOS assembly factors
Mean nuclear PBS in Alaska
Mean nuclear PBS in British Columbia
tmem126a
aifm1
Other genes
Top 5% in both
Correlation between nuclear PBS and mtDNA PBS
cox16
tmem177
cox10
pnkd
lyrm7
NS
p < 0.05
B. Mitochondrial aminoacyl-tRNA synthetases
lars2
pars2
AK top 5% only
BC top 5% only
Other genes
cars2
qrsl1
rars2
gatc
vars2
C. Mitochondrial ribosomal proteins
mrpl23
mrpl13
mrps21
mrpl18
dap3
mrpl51
mrpl16
mrpl14
mrps34
mrps26
mrpl24
mrps2
mrpl27
D. Cytosolic aminoacyl-tRNA synthetases
hars
gars
aimp
kars
dars
eef1e1
E. Cytosolic ribosomal proteins
rpl18
rpl7
uba52
rpl26
rplp0
rps17
rps8a
rpl39
rps6
rpl27

**Figure S7. Concordance between nuclear and mitochondrial PBS across additional mitochondrial-associated and cytosolic gene classes.** Left-hand panels compare mean nuclear population branch statistics (PBS) between British Columbia and Alaska; dashed diagonal lines indicate equal mean differentiation between regions. Right-hand panels show gene-wise correlations between nuclear PBS and mitochondrial PBS across freshwater populations, with statistical significance represented as $-\log_{10}(P)$. Triangles indicate nominally significant correlations ($P < 0.05$), whereas circles indicate nonsignificant correlations. Gene sets comprise (A) OXPHOS assembly factors, (B) mitochondrial aminoacyl-tRNA synthetases, (C) mitochondrial ribosomal proteins, (D) cytosolic aminoacyl-tRNA synthetases, and (E) cytosolic ribosomal proteins.

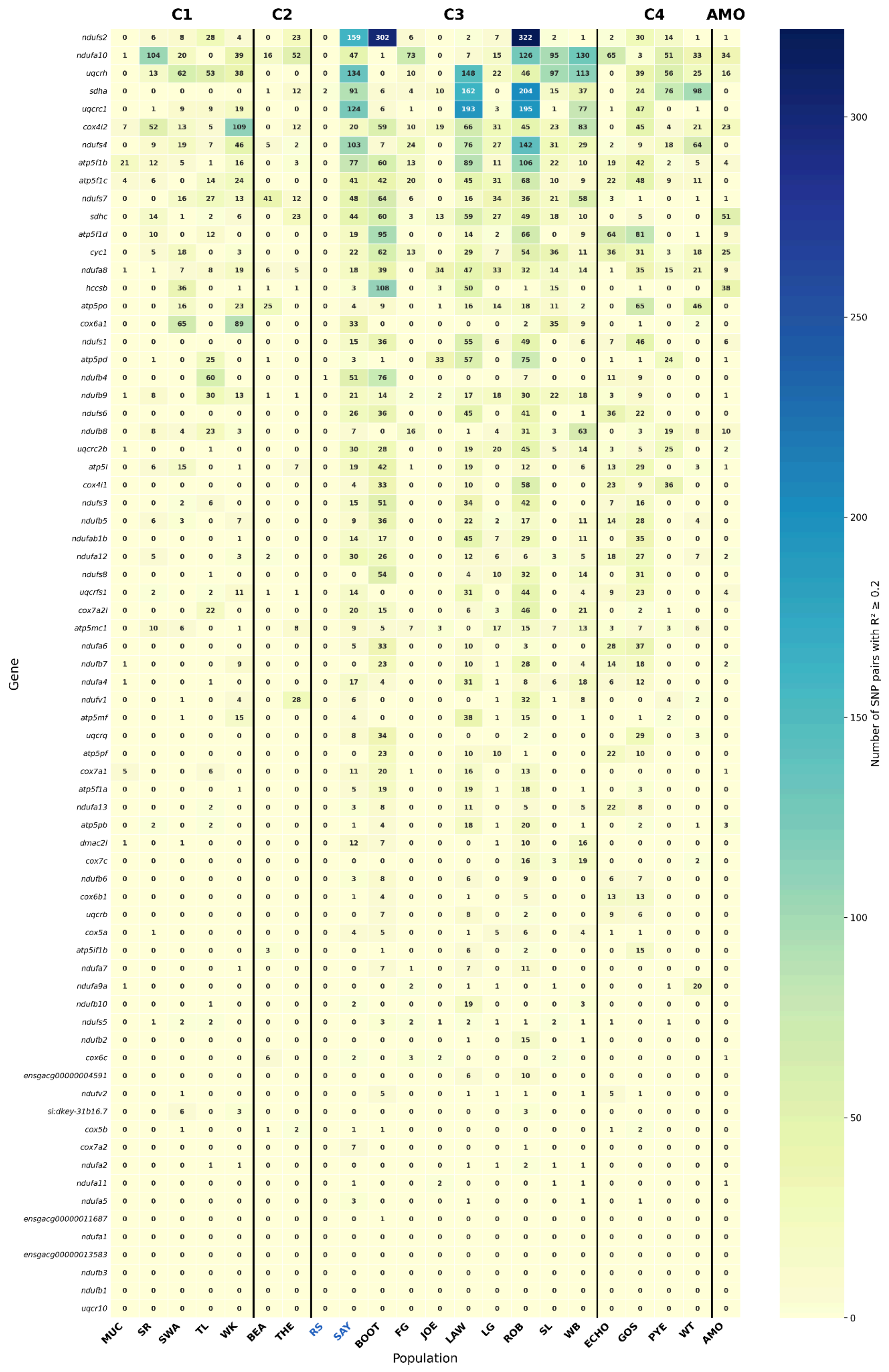

C1
C2
C3
C4
AMO
Gene
ndufs2
ndufa10
uqcrh
sdha
uqcrc1
cox4i2
ndufs4
atp5f1b
atp5f1c
ndufs7
sdhc
atp5f1d
cyc1
ndufa8
hccsb
atp5po
cox6a1
ndufs1
atp5pd
ndufb4
ndufb9
ndufs6
ndufb8
uqcrc2b
atp5l
cox4i1
ndufs3
ndufb5
ndufab1b
ndufa12
ndufs8
uqcrfs1
cox7a2l
atp5mc1
ndufa6
ndufb7
ndufa4
ndufv1
atp5mf
uqcrq
atp5pf
cox7a1
atp5f1a
ndufa13
atp5pb
dmac2l
cox7c
ndufb6
cox6b1
uqcrb
cox5a
atp5if1b
ndufa7
ndufa9a
ndufb10
ndufs5
ndufb2
cox6c
ensgacg00000004591
ndufv2
si:dkey-31b16.7
cox5b
cox7a2
ndufa2
ndufa11
ndufa5
ensgacg00000011687
ndufa1
ensgacg00000013583
ndufb3
ndufb1
uqcr10
MUC
SR
SWA
TL
WK
BEA
THE
RS
SAY
BOOT
FG
JOE
LAW
LG
ROB
SL
WB
ECHO
GOS
PYE
WT
AMO
Population
Number of SNP pairs with R² ≥ 0.2
300
250
200
150
100
50
0

**Figure S8. Linkage disequilibrium structure across nuclear OXPHOS genes and freshwater populations.** Heatmap shows the number of within-gene SNP pairs with $R^2 \geq 0.2$ in each population, as estimated from Pool-seq data using LDx. Established freshwater populations with mitochondrial-lineage assignments are grouped by lineage (C1–C4), and the regional marine reference populations RS and SAY are labeled in blue. AMO was included as an established British Columbia freshwater population but is shown separately because it was not assigned to one of the four mitochondrial lineages. The five recently colonized populations were excluded. These estimates were used to identify linked SNP groups for pruning before SNP-level association analyses.

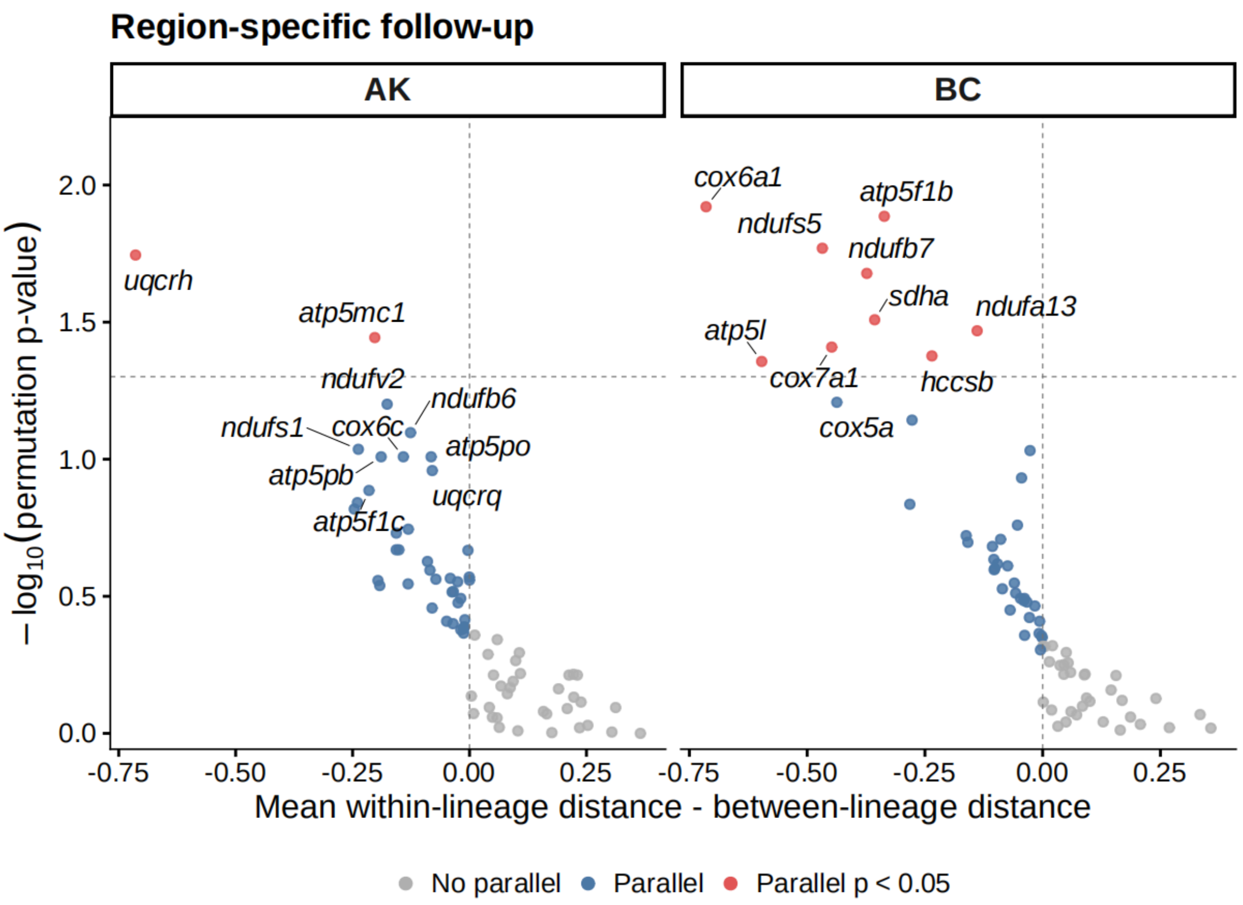


**Figure S9. Region-specific mitochondrial-lineage similarity in nuclear OXPHOS allele-frequency trajectories.** Gene-level comparisons were performed separately in Alaska and British Columbia using LD-pruned ΔAF profiles. Negative x-axis values indicate greater similarity among freshwater populations sharing the same mitochondrial lineage than among populations from different lineages. The vertical dashed line indicates equal within- and between-lineage distances, and the horizontal dashed line indicates permutation $P = 0.05$. Red points identify permutation-supported genes. Analyses excluded AMO and did not apply an SNP effect-size threshold.
